# EpyNN: Educational python for Neural Networks

**DOI:** 10.1101/2021.12.06.470764

**Authors:** Florian Malard, Laura Danner, Emilie Rouzies, Jesse G Meyer, Ewen Lescop, Stéphanie Olivier-Van Stichelen

## Abstract

**Summary:** Artificial Neural Networks (ANNs) have achieved unequaled performance for numerous problems in many areas of Science, Business, Public Policy, and more. While experts are familiar with performance-oriented software and underlying theory, ANNs are difficult to comprehend for non-experts because it requires skills in programming, background in mathematics and knowledge of terminology and concepts. In this work, we release EpyNN, an educational python resource meant for a public willing to understand key concepts and practical implementation of scalable ANN architectures from concise, homogeneous and idiomatic source code. EpyNN contains an educational Application Programming Interface (API), educational workflows from data preparation to ANN training and a documentation website setting side-by-side code, mathematics, graphical representation and text to facilitate learning and provide teaching material. Overall, EpyNN provides basics for python-fluent individuals who wish to learn, teach or develop from scratch.

**Availability:** EpyNN documentation is available at https://epynn.net and repository can be retrieved from https://github.com/synthaze/epynn.

**Contact:** Stéphanie Olivier-Van-Stichelen.

**Supplementary Information:** Supplementary files and listings.

## 1 Introduction

Approaches based on Artificial Neural Networks (ANNs) are implemented to solve problems in many fields with strong implications for topics of public interest such as life sciences and medicine. Therein, highly promising applications of ANNs may include drug discovery [1], gene interaction and disease prediction [2] as well as protein structure prediction [3]. However, the day-to-day integration of ANNs in data analysis workflows remains mostly tied to the expert community, well at ease with ANN theory and Application Programming Interface (API) available from performance-oriented, widespread software [4, 5, 6, 7]. Limitations in programming skills, background in mathematics or terminology and concepts likely explain why deep learning is not standard in every lab dealing with large sets of data.

Non-experts still have opportunities to become familiar with ANNs. The proficient programmer can take advantage of performance-oriented high-level APIs to achieve state-of-the-art results. This, however, requires using ANNs as black-boxes with no understanding of the inner workings. An expert programmer may find readable source codes on sharing platforms but will face highly heterogeneous contents. On the other hand, people with limited programming skills can rely on the extremely rich web documentation including mainstream articles, programming and/or mathematics oriented posts and notebooks. However, while this documentation contains useful items, they do not provide functional codes and are time-consuming.

To cope with the before-mentioned limitations, we release an integrated Educational python resource for Neural Networks (EpyNN) providing homogeneous implementations of diverse ANNs architectures. The EpyNN education-oriented API is written concisely and exhaustively commented to facilitate learning and use as teaching material. The repository contains examples of workflows from data preparation to ANNs training and prediction. EpyNN comes along with a documentation website which provides side-by-side code, mathematics and graphical representations along with standard package documentation. Overall, EpyNN is directed toward non-experts of the field who wish to learn, teach or develop from scratch.

## 2 Implementation

EpyNN is written in Python (3.7.1) [8] and computational flows are written in pure NumPy, the worldwide standard for array programming [9, 10]. Educational examples for data preparation and training of ANNs using EpyNN are provided as regular Python code and Jupyter notebook [11]. The EpyNN documentation website available at https://epynn.net was built with Sphinx [12] using a theme provided by “Read the Docs” [13] and runs with Apache2 [14] over HTTP with SSL/TLS [15] on GNU/Linux Debian 10 (Buster) [16]. EpyNN is licensed under the GNU General Public License v3.0 [17] and is compatible with GNU/Linux, MacOS and Windows.

## 3 Features

### 3.1 Educational API

ANN models are built by stacking layers with distinct architectures (Figure 1). EpyNN implements a set of major layers extensively documented at https://epynn.net. This includes the following layers: *Fully connected* (Dense) [18, 19], *Recurrent Neural Network* (RNN) [20], *Gated Recurrent Unit* (GRU) [21], *Long Short-Term Memory* (LSTM) [22], *Convolution 2D* and *Pooling* layers [23, 24]. EpyNN also provides *Dropout* [25] and *Flatten* layers for network regularization and data reshaping, respectively, as well as a *Template* pass-through layer. Finally, the input *Embedding* layer supports data normalization, one-hot encoding and mini-batches preparation for Stochastic Gradient Descent (SGD). ANN can be created using the *EpyNN* model provided with a list of layers (See https://epynn.net/epynn). Hyperparameters can be set globally for all layers or locally for each layer. Supported activation functions include the Rectifier Linear Unit (ReLU) and Leaky ReLU, Exponential Linear Unit (ELU) and logistic functions sigmoid, hyperbolic tangent and softmax (See https://epynn.net/activation). Supported loss functions include Mean Squared Error (MSE), Mean Absolute Error (MAE), Mean Squared Logarithmic Error (MSLE), Binary Cross-Entropy (BCE) and Categorical Cross-Entropy (CCE) (See https://epynn.net/loss). Evaluation metrics include accuracy, precision, recall, specificity and F-Score.

**Figure 1:**
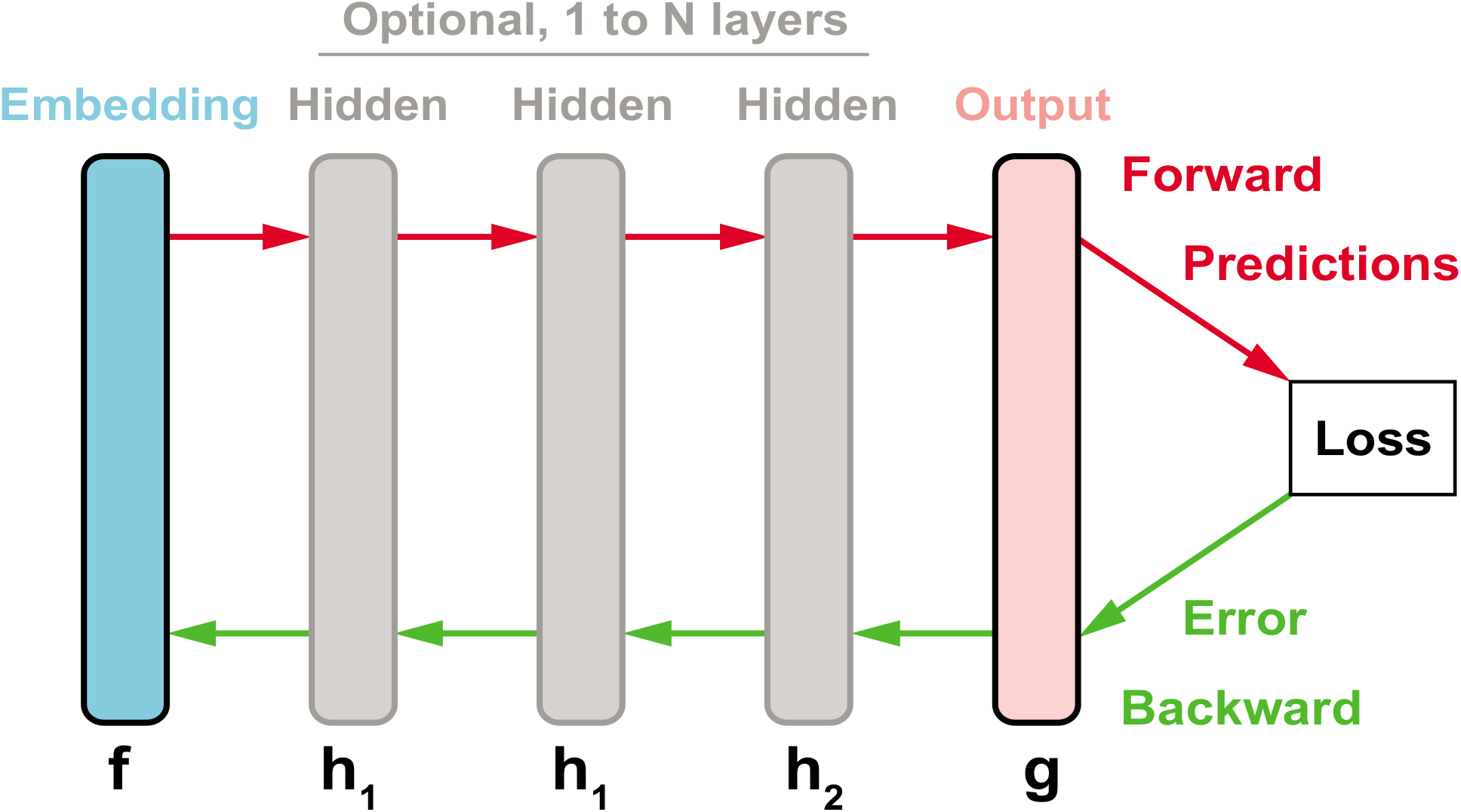
ANN model and layers. Here is depicted a set of five layers, equivalent to a set of five functions {*f, h*_1_, *h*_1_, *h*_2_, *g*}. This set contains a subset of four distinct layer architectures {*f, h*_1_, *h*_2_, *g*} because two layers are defined by the same function *h*_1_. The ANN model featuring the set of five stacked layers is the composite function (*g* ∘ *h*_2_ ∘ *h*_1_ ∘ *h*_1_ ∘ *f*) which takes input and returns predictions (forward, red) further compared to target values through the derivative of a loss function (e.g., Mean Squared Error). The error gradient is propagated backward through each layer and, if applicable, trainable parameters are updated accordingly (backward, green). See https://epynn.net/epynn for more details.

### 3.2 Educational Python

One layer is defined inside one directory containing four files for model definition, forward propagation, backward propagation and parameters-related functions (See https://github.com/synthaze/epynn/tree/main/epynn). Sources strictly layer mathematical definitions and therefore they are accurate and concise with definition ranging from 66 to 269 lines of codes for the *Template* and LSTM layers, respectively. The ANN model *EpyNN* and other modules for activation, loss functions, metrics and more were written following the same guidelines. Importantly, sources are extensively commented with an average of 0.35 line of comments for each line of code (Table 1). Overall, native implementations are written in idiomatic Python/NumPy with a strong focus on homogeneity across the whole package.

**Table 1:**
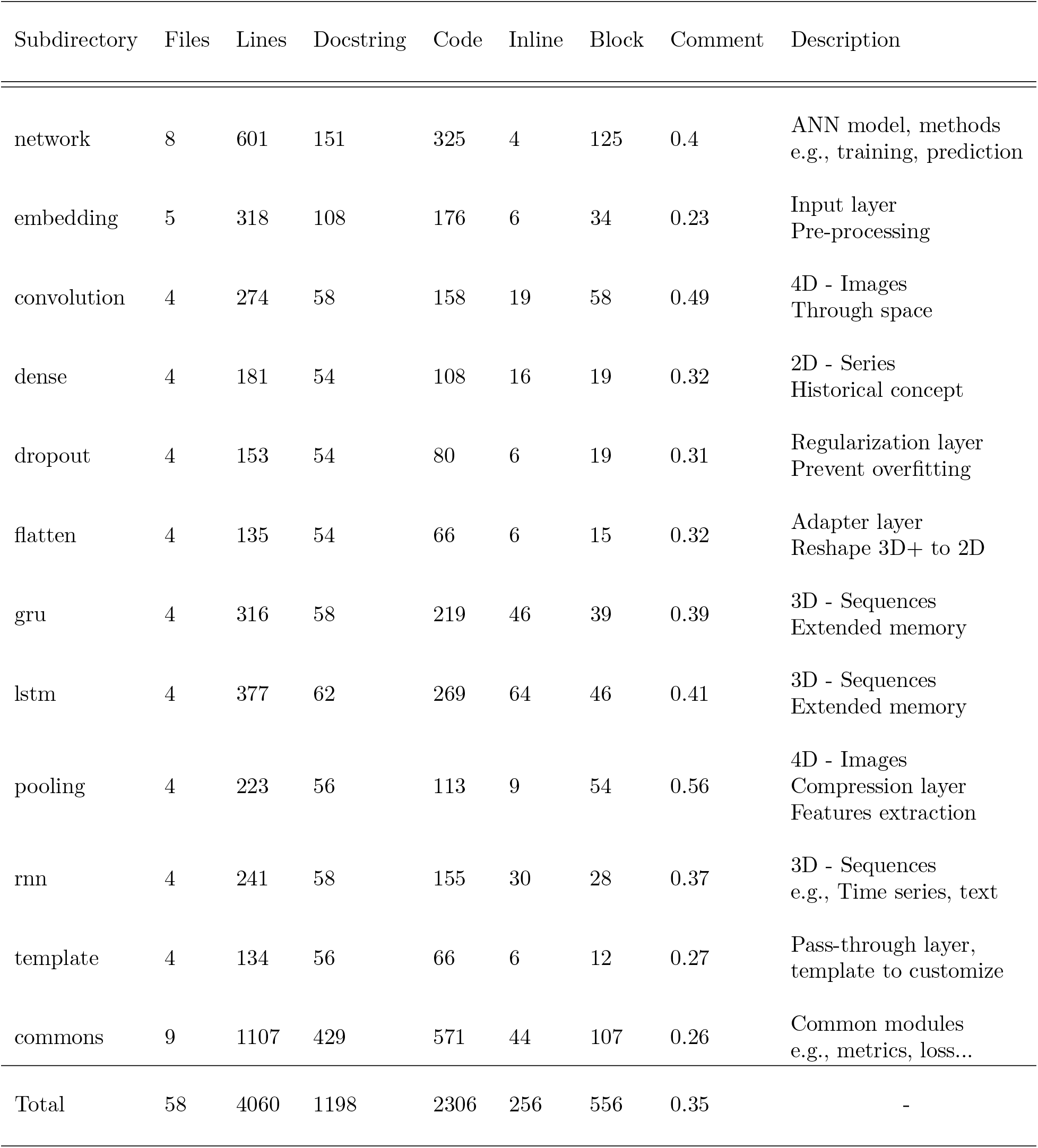
EpyNN *epynn* library tree - Description and source code statistics. The educational library module in EpyNN repository corresponds to the *epynn* directory. Each subdirectory except *network* and *commons* implement a specific layer architecture. RNN, LSTM and GRU are so-called recurrent layers. Files: Number of files in each subdirectory. Lines: Total number of lines in files,excluding blank lines. Docstring: Lines accounting for documentation strings or “global” code comments. Code: Lines accounting for executable python code. Inline, Block: Lines accounting for “local” code comments. Comment: Is equal to 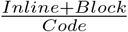 or lines of “local” code comments for one line of executable code. Review the *epynnlive* directory at https://github.com/synthaze/epynn/tree/main/epynn.

| Subdirectory | Files | Lines | Docstring | Code | Inline | Block | Comment | Description |
| --- | --- | --- | --- | --- | --- | --- | --- | --- |
| network | 8 | 601 | 151 | 325 | 4 | 125 | 0.4 | ANN model, methods<br>e.g., training, prediction |
| embedding | 5 | 318 | 108 | 176 | 6 | 34 | 0.23 | Input layer<br>Pre-processing |
| convolution | 4 | 274 | 58 | 158 | 19 | 58 | 0.49 | 4D - Images<br>Through space |
| dense | 4 | 181 | 54 | 108 | 16 | 19 | 0.32 | 2D - Series<br>Historical concept |
| dropout | 4 | 153 | 54 | 80 | 6 | 19 | 0.31 | Regularization layer<br>Prevent overfitting |
| flatten | 4 | 135 | 54 | 66 | 6 | 15 | 0.32 | Adapter layer<br>Reshape 3D+ to 2D |
| gru | 4 | 316 | 58 | 219 | 46 | 39 | 0.39 | 3D - Sequences<br>Extended memory |
| lstm | 4 | 377 | 62 | 269 | 64 | 46 | 0.41 | 3D - Sequences<br>Extended memory |
| pooling | 4 | 223 | 56 | 113 | 9 | 54 | 0.56 | 4D - Images<br>Compression layer<br>Features extraction |
| rnn | 4 | 241 | 58 | 155 | 30 | 28 | 0.37 | 3D - Sequences<br>e.g., Time series, text |
| template | 4 | 134 | 56 | 66 | 6 | 12 | 0.27 | Pass-through layer,<br>template to customize |
| commons | 9 | 1107 | 429 | 571 | 44 | 107 | 0.26 | Common modules<br>e.g., metrics, loss... |
| Total | 58 | 4060 | 1198 | 2306 | 256 | 556 | 0.35 | - |

### 3.3 Educational Workflows

We provide 14 educational workflows on data type, structure and preparation along with principles to design and train ANN, both in regular Python and Jupyter notebooks formats (Table 2). In the notebooks, we introduced educational features of EpyNN’s API. Among others, this includes exhaustive initialization logs reporting on layers’ dimensions and shapes for input, output and processing intermediates. In addition, layers in EpyNN share the same cache structure, which makes easy content manipulation out of built-in procedures and purposes. To facilitate monitoring, we implemented colorful logs during model training along with automatic plot facilities upon completion.

**Table 2:** EpyNN *epynnlive* examples tree - Description and source code statistics. Educational workflows for data preparation and model training are located inside the *epynnlive* directory in the EpyNN repository. Each subdirectory contains 3 python files *prepare_dataset.py, train.py* and *set-tings.py* and Jupyter notebooks versions *prepare_dataset.ipynb* and *train.ipynb* not accounted here. In descrption, CNN refers to Convolutional Neural Network. Lines: Total number of lines in files, excluding blank lines. Docstring: Lines accounting for documentation strings or “global” code comments. Code: Lines accounting for executable python code. Block: Lines accounting for “local” code comments. Comment: Is equal to 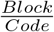 or lines of “local” code comments for one line of executable code. Models: Number of distinct ANN configuration reviewed in *train.py*. Review the *epynnlive* directory at https://github.com/synthaze/epynn/tree/main/epynnlive and Jupyter notebooks for data preparation at https://epynn.net/nbdata and model training at https://epynn.net/nbtraining.

| Subdirectory | Files | Lines | Docstring | Code | Block | Comment | Models | Description |
| --- | --- | --- | --- | --- | --- | --- | --- | --- |
| dummy_boolean | 3 | 122 | 26 | 64 | 32 | 0.52 | 1 | Python basics, Boolean, Perceptron |
| dummy_image | 3 | 174 | 32 | 100 | 42 | 0.42 | 2 | Image definition and generation, CNN |
| dummy_string | 3 | 169 | 26 | 106 | 37 | 0.35 | 4 | String data type, one-hot encoding, recurrent layers |
| dummy_time | 3 | 196 | 34 | 119 | 43 | 0.36 | 4 | Time series, signal detection, recurrent layers |
| author_music | 3 | 209 | 23 | 138 | 48 | 0.35 | 3 | WAV audio files manipulation, recurrent layers |
| captcha_mnist | 3 | 148 | 13 | 104 | 31 | 0.3 | 2 | IDX format, encoded image, CNN |
| ptm_protein | 3 | 178 | 13 | 127 | 38 | 0.3 | 4 | O-GlcNAcylated peptides [26, 27], recurrent layers |
| Total | 21 | 1196 | 167 | 758 | 271 | 0.36 | 20 | - |

### 3.4 Educational Website

https://epynn.net was designed to provide integrative educational material, traditional python package documentation and Jupyter notebooks. Therein, systematic descriptions put side-by-side code, mathematical notation, graphical representation and succinct text-based explanations for most object in EpyNN (Figure 2). Because ANNs may be difficult to comprehend for non-expert, we consider this multi-levels approach is key to facilitate understanding with regards to the diversity of users and backgrounds. Accordingly, we expect more individuals to find an anchor point and therefore evolve toward a global understanding. Overall, we implemented https://epynn.net to be an easy learning material and teaching support.

**Figure 2:**
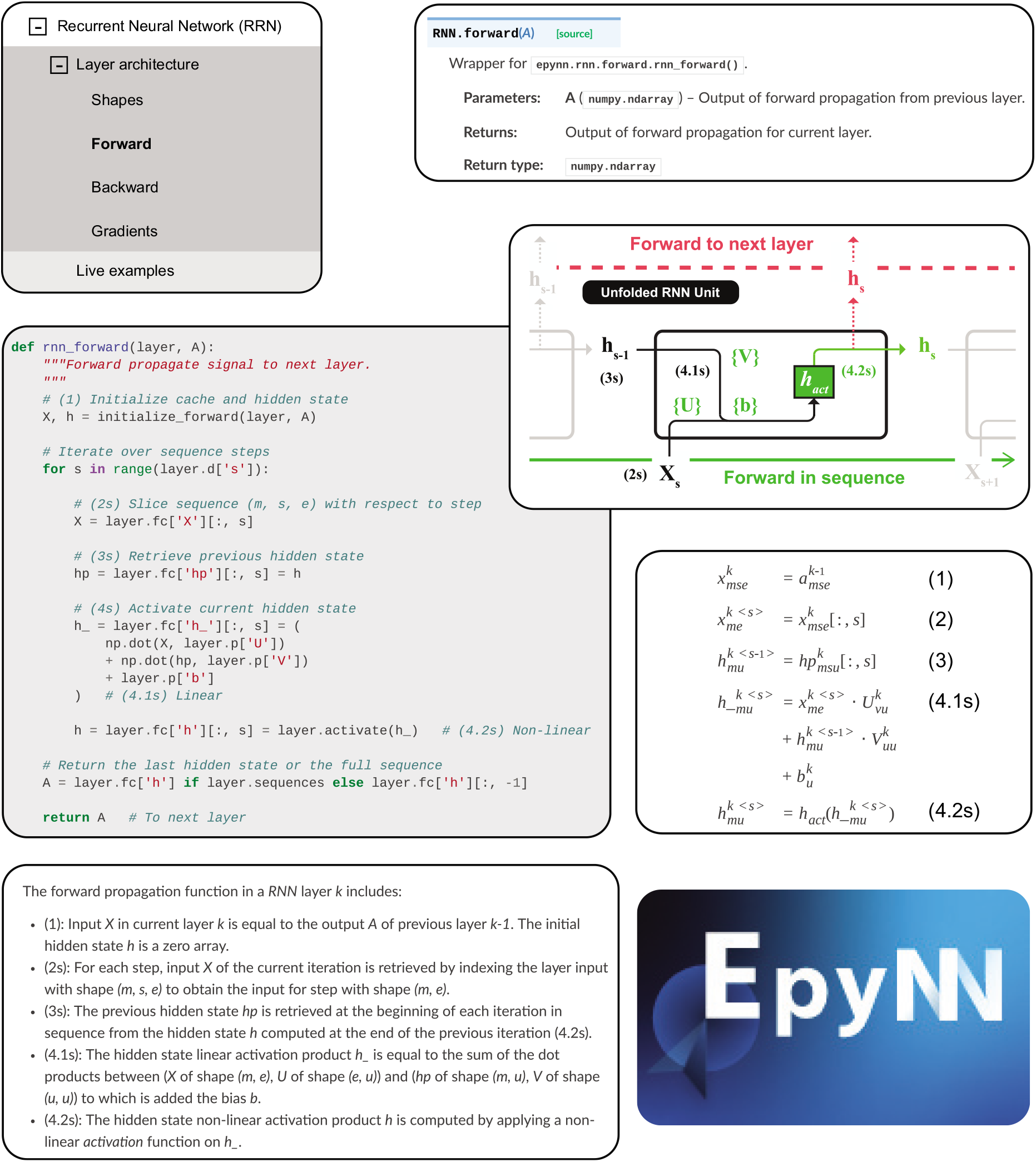
EpyNN https://epynn.net educational website - Example of integrated documentation. The EpyNN educational API is documented with systematic descriptions involving classic API documentation and corresponding source codes explained line-by-line via direct translation into diagrams, mathematics and text. This scheme is shown for the RNN layer and is extracted from https://epynn.net/RNN. Objects and methods/functions related to network and layer models or activation and loss functions, among others, were documented following this scheme. See https://epynn.net/glossary#notations for conventions about notations.

## 4 Related work

EpyNN should not be confounded in the crowded space of deep-learning libraries that attempt to provide a high-level API that is easy to use. Such popular deep-learning libraries are generally developed for performance in production environment and not for educational purposes (*e.g*., Tensorflow [5], Keras [4], Torch [6], Fastai [7], and others). The main and crucial difference is simple: the source code of high-level APIs in production-focused deep-learning libraries is not written to be used as a teaching or learning material. This means that it turns extremely difficult to locate and understand the algorithms explaining the inner working of ANNs in the source code of such popular, production-focused libraries (Listing S1, S2).

Still, EpyNN is highly suited to teach and learn the backbone of other popular, production-focused libraries. EpyNN computational schemes were validated by direct comparison with Tensorflow/Keras [4] in 264 distinct configurations (File S1). While providing identical results for identical configurations, the *tensorflow/python* directory contains 1291 python files for a total of 333 527 lines of codes (File S2). By contrast, the *EpyNN/epynn* directory contains 58 files for a total of 2317 lines of code. Therefore, EpyNN provides the opportunity to read and understand every line of code behind its educational API which may remain highly similar in use compared with other libraries, including Tensorflow/Keras [4].

## 5 Context of Use

The main goal of EpyNN is to provide an integrated environment allowing to understand the inner working of ANNs trained with backpropagation. EpyNN brings together theory and practice through its lightweight API which relies on sources that are written to be read and modified by users. This in conjunction with an educational website that describes the functional code line by line with diagrams, mathematics and text. Said differently, there is virtually no abstraction between the source code of EpyNN and the integrated documentation.

Typically, one user would use EpyNN by setting up the following environment: one text editor session to review and customize the source code of EpyNN educational API, another session to write the executable script for ANN design and training, a web session at https://epynn.net to take advantage of the extended documentation and finally a terminal session to proceed with ANN training and further operations (Figure 3). Users may work on a local copy of EpyNN educational API sources to design exercises, to customize and compare variants of existing architectures or even to implement new architecture via the *Template* layer (See https://epynn.net/epynnlayers#template-layer).

**Figure 3:**
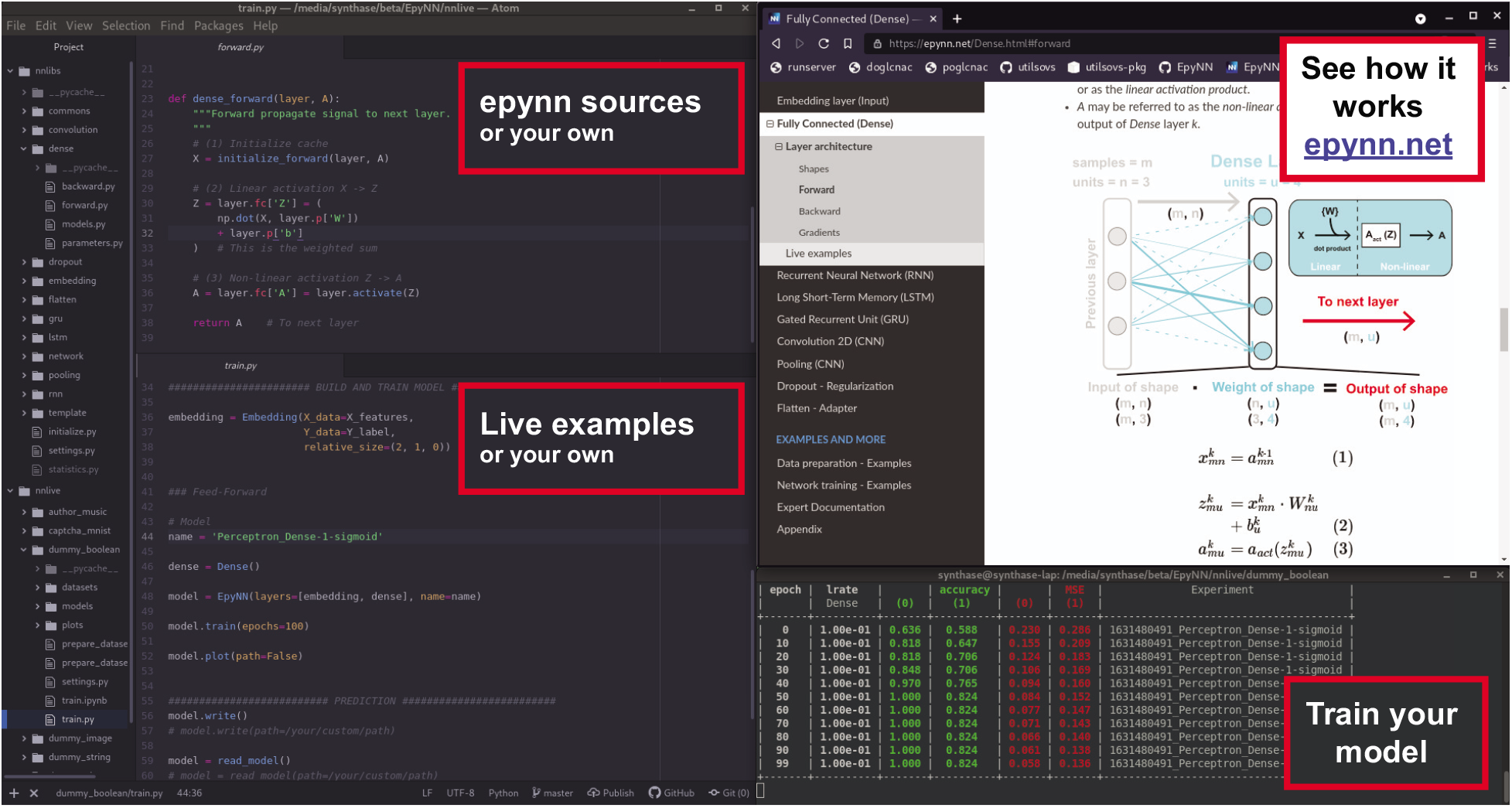
How to use EpyNN - Suggestion of interface. The interface is made of text editor sessions for EpyNN *epynn* library (left, top) and live examples (left, bottom), EpyNN https://epynn.net educational website (right, top) and a terminal session to run EpyNN (right, bottom). EpyNN sources can refer to the root module files or a working copy in the current directory. Note that new layer architectures can be implemented on-the-fly from existing and template layers. See https://epynn.net/epynnlayers and https://epynn.net/quickstart#how-to-use-epynn for more documentation.

## 6 Conclusion

We developed EpyNN, an education-oriented python resource aiming to encourage practice, understanding and adoption of ANNs by non-experts researchers and beyond. EpyNN features an educational API, concise python sources, educational workflows and a rich educational website. Because EpyNN is easy to customize, we anticipate and encourage deposition of additional content from push request at https://github.com/synthaze/epynn.

## Acknowledgments

We thank Axelle Malard (https://axellemalard.com), artistic director and graphic designer who kindly offered to provide EpyNN logos and favicons. We thank Christelle Gloor (D-INFK, ETH Zurich) for useful discussions during the development of EpyNN. This work was supported by the Medical College of Wisconsin and the National Institute of Health (R00 HD087430).

## Supplementary Material

### Supplementary files

#### Supplementary file S1

FileS1.zip: Contains executable python codes used for validation of EpyNN against Tensorflow/Keras using tensorflow==2.3.0 pip package. See README for details. Note that a summary of results is available at https://epynn.net/index#is-epynn-reliable.

#### Supplementary file S2

FileS2.zip: XLSX file summary and python script used to count the number of files and lines of code within tensorflow/python directory from tensorflow==2.3.0 pip package. By lines of code we mean all but not blank lines, docstrings and block comments.

### Supplementary listings

#### Supplementary listing S1

**Listing S1:**
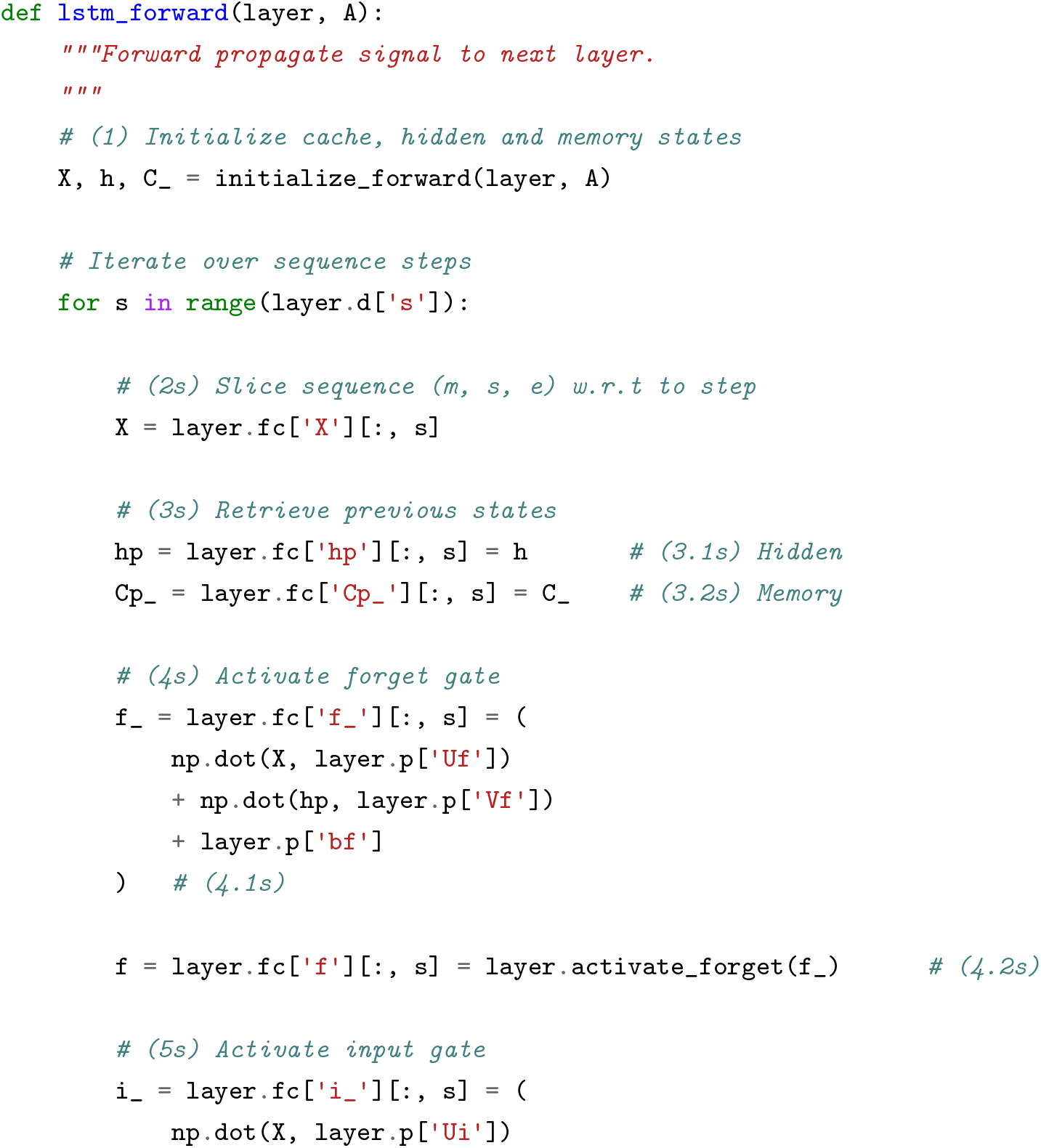
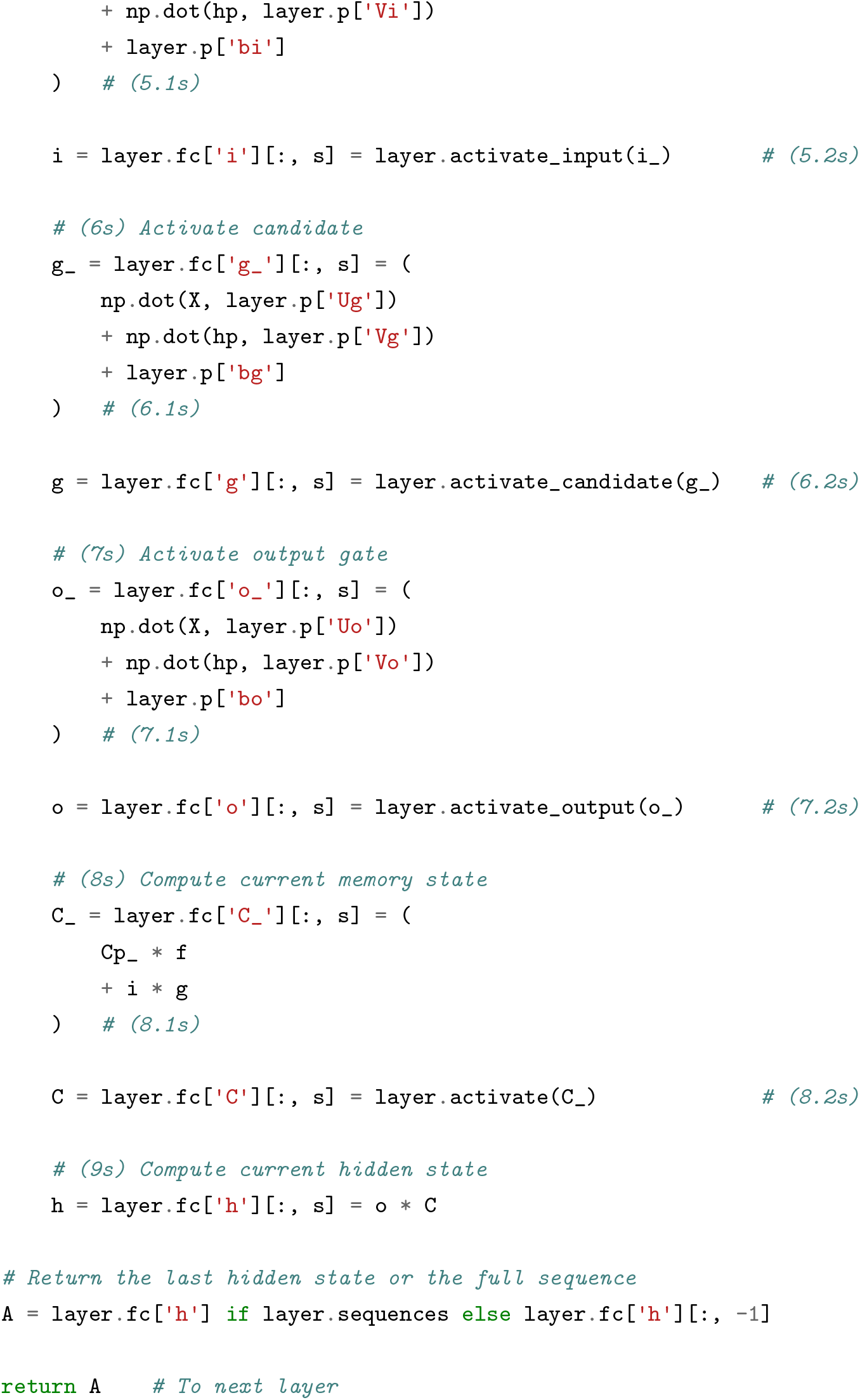
EpyNN sources for *epynn.lstm.forward.lstm_forward()*. The function describes the forward propagation algorithm for the LSTM layer in EpyNN. This function is easy to find in the source of EpyNN because it is simply located within the *epynn.lstm.forward* module which contains a total of two functions for less than hundred lines of code. Moreover, it is easy to understand because written in idiomatic Python/NumPy and exhaustively commented. Finally, extended documentation on this code is available at https://epynn.net/LSTM#forward. See https://epynn.net/glossary#notations for conventions about notations.

#### Supplementary listing S2

**Listing S2:**
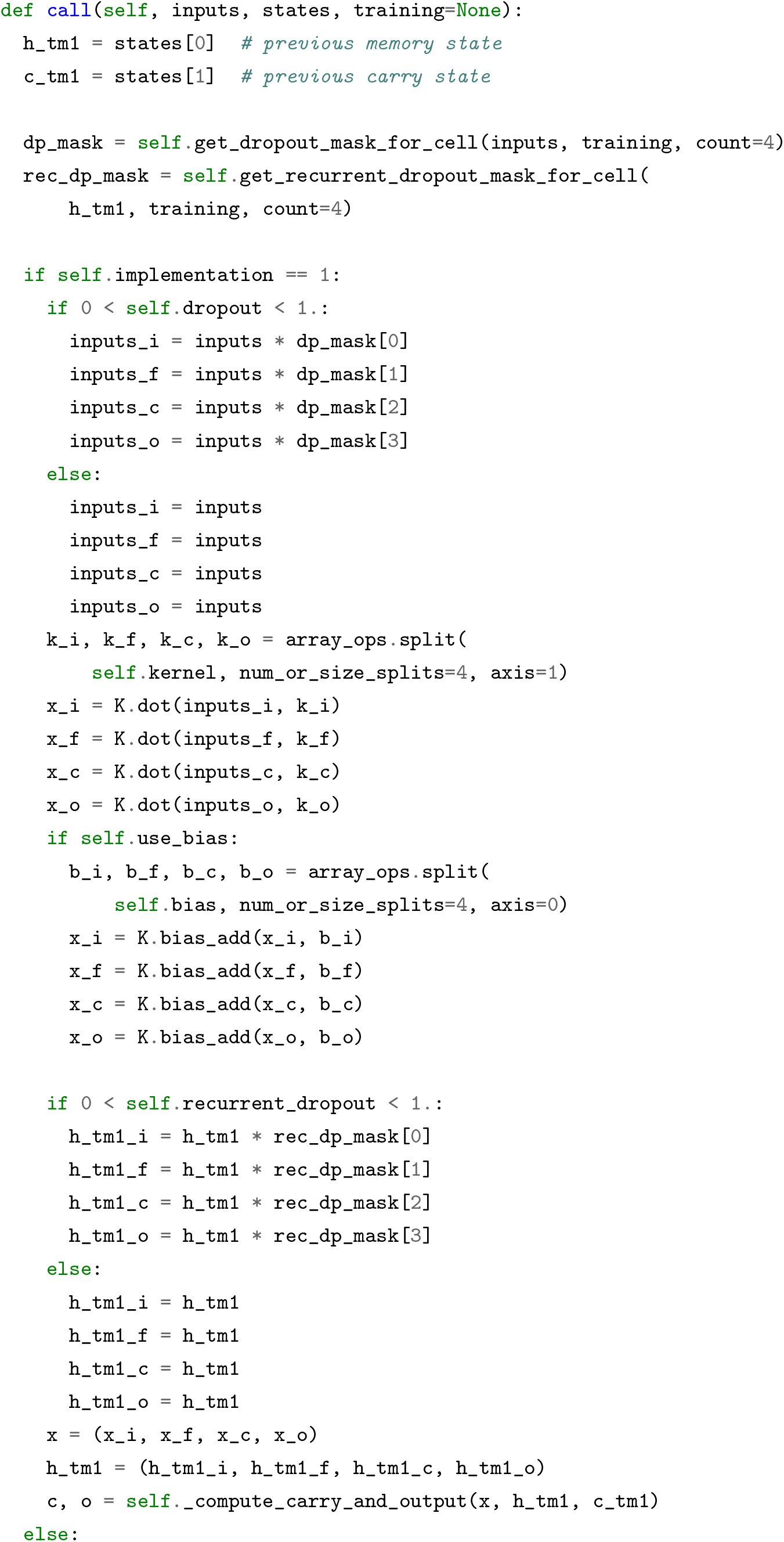
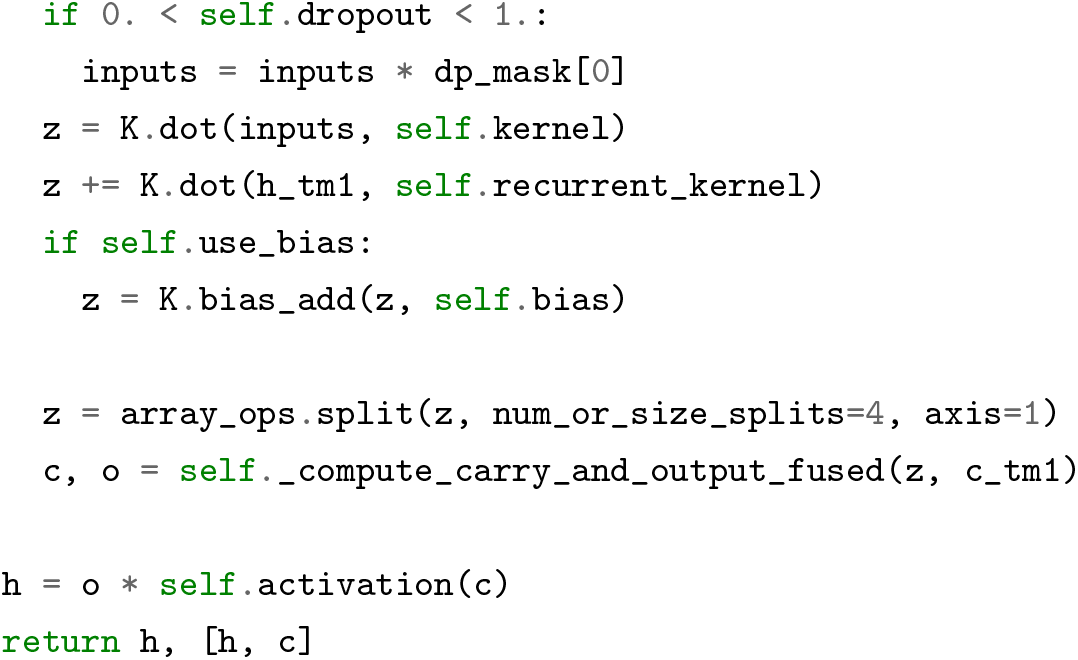
Tensorflow/Keras sources for *tensorflow.python.keras.layers.recurrent.LSTMCell.call()* The method describes the forward propagation algorithm for the LSTM cell in Tesorflow/Keras. This method is not easy to find in the source of Tensorflow/Keras because of the number of recursion levels down to *tensorflow.python.keras.layers.recurrent* module which contains 34 objects, 200 methods and 1 766 lines of code. In addition, there is no block or inline comment in this code.

## Notes

https://EpyNN.net

### Competing Interest Statement

The authors have declared no competing interest.

