## Supplementary material for "EpyNN: Educational python for Neural Networks": File S1: plot.py

```
import matplotlib.pyplot as plt
import numpy as np


rmsd_log = {k: [0, 0] for k in [-8, -7, -6, -5, -4, -4, -2, -1, 0]}

with open('autotest_proc.dat', 'r') as f:
    for line in f:
        kloss = float(line.split(',')[0])
        preds = float(line.split()[1])
        for power in rmsd_log:
            if kloss < 10**power:
                rmsd_log[power][0] += 1
                break
        for power in rmsd_log:
            if preds < 10**power:
                rmsd_log[power][1] += 1
                break

x = np.array(list(rmsd_log.keys()))

dL = np.array([x[0] for x in rmsd_log.values()])

dP = np.array([x[1] for x in rmsd_log.values()])

fig, axs = plt.subplots(1, 2)

axs[0].bar(x, dL)
axs[0].set_xticks([-8, -7, -6, -5, -4, -3, -2, -1, 0])
axs[0].set_yticks([0, 50, 100, 150, 200, 250])
axs[0].set_title('RMSD - Loss')
axs[0].set_xlabel('RMSD < 10**X')
axs[0].set_ylabel('Number of Experiments')

axs[1].bar(x, dP)
axs[1].set_xticks([-8, -7, -6, -5, -4, -3, -2, -1, 0])
axs[1].set_yticks([0, 50, 100, 150, 200, 250])
axs[1].set_title('RMSD - Probabilities')
axs[1].set_xlabel('RMSD < 10**X')
axs[1].set_ylabel('Number of Experiments')

plt.tight_layout()
plt.savefig('keras_epynn.svg')

print(rmsd_log)
```
