## Supplementary material for "EpyNN: Educational python for Neural Networks": File S1: autotest_Conv_Pool_Dense.py

```
"""Automatic testing.
"""
import os

### Conv_Pool_Dense.py

sigmoid_configs = {
    'conv0_filters': [4, 4, 4, 4, 4, 4, 4, 4, 4],
    'conv0_in': [False, False, False, False, False, False, True, True, True],

    'pool0_in': [False, False, False, False, False, False, True, True, True],

    'conv1_filters': [2, 4, 8, 2, 4, 8, 2, 4, 8],
    'conv1_in': [True, True, True, True, True, True, True, True, True],

    'pool1_in': [False, False, False, True, True, True, True, True, True],

    'dense_units': [1, 1, 1, 2, 2, 2, 4, 4, 4],
    'dense_activation': ['sigmoid', 'sigmoid', 'sigmoid', 'sigmoid', 'sigmoid', 'sigmoid', 'sigmoid', 'sigmoid', 'sigmoid'],
    'dense_trainable': [False, False, False, False, False, False, False, False, False],

    'LOSS': ['MSE', 'MAE', 'MSLE', 'MSE', 'MAE', 'MSLE', 'MSE', 'MAE', 'MSLE'],
}


tanh_configs = {
    'conv0_filters': [4, 4, 4, 4, 4, 4],
    'conv0_in': [False, False, False, False, True, True],

    'pool0_in': [False, False, False, False, True, True],

    'conv1_filters': [2, 4, 8, 2, 4, 8],
    'conv1_in': [True, True, True, True, True, True],

    'pool1_in': [False, False, False, True, True, True],

    'dense_units': [1, 1, 2, 2, 4, 4],
    'dense_activation': ['tanh', 'tanh', 'tanh', 'tanh', 'tanh', 'tanh',],
    'dense_trainable': [True, True, True, True, True, True],

    'LOSS': ['MSE', 'MAE', 'MSE', 'MAE', 'MSE', 'MAE'],
}


softmax_configs = {
    'conv0_filters': [4, 4, 4, 4, 4],
    'conv0_in': [True, True, True, True, True],

    'pool0_in': [True, True, True, True, True],

    'conv1_filters': [1, 2, 4, 8, 16],
    'conv1_in': [True, True, True, True, True],

    'pool1_in': [True, True, True, True, True],

    'dense_units': [2, 2, 2, 2, 2],
    'dense_activation': ['softmax', 'softmax', 'softmax', 'softmax', 'softmax'],
    'dense_trainable': [True, True, True, True, True],

    'LOSS': ['MSE', 'MAE', 'MSLE', 'BCE', 'CCE'],
}


bis_softmax_configs = {
    'conv0_filters': [4, 4, 4, 4, 4],
    'conv0_in': [True, True, True, True, True],

    'pool0_in': [True, True, True, True, True],

    'conv1_filters': [1, 2, 4, 8, 16],
    'conv1_in': [True, True, True, True, True],

    'pool1_in': [True, True, True, True, True],

    'dense_units': [4, 4, 4, 4, 4],
    'dense_activation': ['softmax', 'softmax', 'softmax', 'softmax', 'softmax'],
    'dense_trainable': [True, True, True, True, True],

    'LOSS': ['MSE', 'MAE', 'MSLE', 'BCE', 'CCE'],
}

for configs in [sigmoid_configs, tanh_configs, softmax_configs, bis_softmax_configs]:

    N_RUNS = len(configs['conv0_filters'])

    for run in range(N_RUNS):
        args = []

        for k, v in configs.items():
            args.append(str(v[run]))
        args = ' '.join(args)

        os.system('python3 Conv_Pool_Dense.py %s' % args)
```
