## Supplementary material for "EpyNN: Educational python for Neural Networks": File S1: autotest_Dense_Dense.py

```
"""Automatic testing.
"""
import os

### Dense_Dense.py

P_sigmoid_configs = {
    'dense0_units': [16, 16, 16, 16, 16, 16, 16, 16, 16],
    'dense0_activation': ['tanh', 'tanh', 'tanh', 'tanh', 'tanh', 'tanh', 'tanh', 'tanh', 'tanh'],
    'dense0_trainable': [False, False, False, False, False, False, False, False, False],
    'dense0_in': [False, False, False, False, False, False, False, False, False],

    'dense1_units': [8, 8, 8, 8, 8, 8, 8, 8, 8],
    'dense1_activation': ['elu', 'elu', 'elu', 'elu', 'elu', 'elu', 'elu', 'elu', 'elu'],
    'dense1_trainable': [False, False, False, False, False, False, False, False, False],
    'dense1_in': [False, False, False, False, False, False, False, False, False],

    'dense_units': [1, 1, 1, 2, 2, 2, 4, 4, 4],
    'dense_activation': ['sigmoid', 'sigmoid', 'sigmoid', 'sigmoid', 'sigmoid', 'sigmoid', 'sigmoid', 'sigmoid', 'sigmoid'],
    'dense_trainable': [True, True, True, True, True, True, True, True, True],

    'LOSS': ['MSE', 'MAE', 'MSLE', 'MSE', 'MAE', 'MSLE', 'MSE', 'MAE', 'MSLE'],
}

P_tanh_configs = {
    'dense0_units': [16, 16, 16, 16, 16, 16],
    'dense0_activation': ['tanh', 'tanh', 'tanh', 'tanh', 'tanh', 'tanh'],
    'dense0_trainable': [False, False, False, False, False, False],
    'dense0_in': [False, False, False, False, False, False],

    'dense1_units': [8, 8, 8, 8, 8, 8],
    'dense1_activation': ['elu', 'elu', 'elu', 'elu', 'elu', 'elu'],
    'dense1_trainable': [False, False, False, False, False, False],
    'dense1_in': [False, False, False, False, False, False],

    'dense_units': [1, 1, 2, 2, 4, 4],
    'dense_activation': ['tanh', 'tanh', 'tanh', 'tanh', 'tanh', 'tanh',],
    'dense_trainable': [True, True, True, True, True, True],

    'LOSS': ['MSE', 'MAE', 'MSE', 'MAE', 'MSE', 'MAE'],
}

P_softmax_configs = {
    'dense0_units': [16, 16, 16, 16, 16],
    'dense0_activation': ['tanh', 'tanh', 'tanh', 'tanh', 'tanh'],
    'dense0_trainable': [False, False, False, False, False],
    'dense0_in': [False, False, False, False, False],

    'dense1_units': [8, 8, 8, 8, 8],
    'dense1_activation': ['elu', 'elu', 'elu', 'elu', 'elu'],
    'dense1_trainable': [False, False, False, False, False],
    'dense1_in': [False, False, False, False, False],

    'dense_units': [2, 2, 2, 2, 2],
    'dense_activation': ['softmax', 'softmax', 'softmax', 'softmax', 'softmax'],
    'dense_trainable': [True, True, True, True, True],

    'LOSS': ['MSE', 'MAE', 'MSLE', 'BCE', 'CCE'],
}

NN_sigmoid_configs = {
    'dense0_units': [16, 16, 16, 16, 16, 16, 16, 16, 16],
    'dense0_activation': ['tanh', 'tanh', 'tanh', 'tanh', 'tanh', 'tanh', 'tanh', 'tanh', 'tanh'],
    'dense0_trainable': [False, False, False, False, False, False, False, False, False],
    'dense0_in': [False, False, False, False, False, False, False, False, False],

    'dense1_units': [8, 8, 8, 8, 8, 8, 8, 8, 8],
    'dense1_activation': ['elu', 'elu', 'elu', 'elu', 'elu', 'elu', 'elu', 'elu', 'elu'],
    'dense1_trainable': [True, True, True, True, True, True, True, True, True],
    'dense1_in': [True, True, True, True, True, True, True, True, True],

    'dense_units': [1, 1, 1, 2, 2, 2, 4, 4, 4],
    'dense_activation': ['sigmoid', 'sigmoid', 'sigmoid', 'sigmoid', 'sigmoid', 'sigmoid', 'sigmoid', 'sigmoid', 'sigmoid'],
    'dense_trainable': [True, True, True, True, True, True, True, True, True],

    'LOSS': ['MSE', 'MAE', 'MSLE', 'MSE', 'MAE', 'MSLE', 'MSE', 'MAE', 'MSLE'],
}

NN_tanh_configs = {
    'dense0_units': [16, 16, 16, 16, 16, 16],
    'dense0_activation': ['tanh', 'tanh', 'tanh', 'tanh', 'tanh', 'tanh'],
    'dense0_trainable': [False, False, False, False, False, False],
    'dense0_in': [False, False, False, False, False, False],

    'dense1_units': [8, 8, 8, 8, 8, 8],
    'dense1_activation': ['elu', 'elu', 'elu', 'elu', 'elu', 'elu'],
    'dense1_trainable': [True, True, True, True, True, True],
    'dense1_in': [True, True, True, True, True, True],

    'dense_units': [1, 1, 2, 2, 4, 4],
    'dense_activation': ['tanh', 'tanh', 'tanh', 'tanh', 'tanh', 'tanh',],
    'dense_trainable': [True, True, True, True, True, True],

    'LOSS': ['MSE', 'MAE', 'MSE', 'MAE', 'MSE', 'MAE'],
}

NN_softmax_configs = {
    'dense0_units': [16, 16, 16, 16, 16],
    'dense0_activation': ['tanh', 'tanh', 'tanh', 'tanh', 'tanh'],
    'dense0_trainable': [False, False, False, False, False],
    'dense0_in': [False, False, False, False, False],

    'dense1_units': [8, 8, 8, 8, 8],
    'dense1_activation': ['elu', 'elu', 'elu', 'elu', 'elu'],
    'dense1_trainable': [True, True, True, True, True],
    'dense1_in': [True, True, True, True, True],

    'dense_units': [2, 2, 2, 2, 2],
    'dense_activation': ['softmax', 'softmax', 'softmax', 'softmax', 'softmax'],
    'dense_trainable': [True, True, True, True, True],

    'LOSS': ['MSE', 'MAE', 'MSLE', 'BCE', 'CCE'],
}


DNN_sigmoid_configs = {
    'dense0_units': [16, 16, 16, 16, 16, 16, 16, 16, 16],
    'dense0_activation': ['tanh', 'tanh', 'tanh', 'tanh', 'tanh', 'tanh', 'tanh', 'tanh', 'tanh'],
    'dense0_trainable': [True, True, True, True, True, True, True, True, True],
    'dense0_in': [True, True, True, True, True, True, True, True, True],

    'dense1_units': [8, 8, 8, 8, 8, 8, 8, 8, 8],
    'dense1_activation': ['elu', 'elu', 'elu', 'elu', 'elu', 'elu', 'elu', 'elu', 'elu'],
    'dense1_trainable': [True, True, True, True, True, True, True, True, True],
    'dense1_in': [True, True, True, True, True, True, True, True, True],

    'dense_units': [1, 1, 1, 2, 2, 2, 4, 4, 4],
    'dense_activation': ['sigmoid', 'sigmoid', 'sigmoid', 'sigmoid', 'sigmoid', 'sigmoid', 'sigmoid', 'sigmoid', 'sigmoid'],
    'dense_trainable': [True, True, True, True, True, True, True, True, True],

    'LOSS': ['MSE', 'MAE', 'MSLE', 'MSE', 'MAE', 'MSLE', 'MSE', 'MAE', 'MSLE'],
}

DNN_tanh_configs = {
    'dense0_units': [16, 16, 16, 16, 16, 16],
    'dense0_activation': ['tanh', 'tanh', 'tanh', 'tanh', 'tanh', 'tanh'],
    'dense0_trainable': [True, True, True, True, True, True],
    'dense0_in': [True, True, True, True, True, True],

    'dense1_units': [8, 8, 8, 8, 8, 8],
    'dense1_activation': ['elu', 'elu', 'elu', 'elu', 'elu', 'elu'],
    'dense1_trainable': [True, True, True, True, True, True],
    'dense1_in': [True, True, True, True, True, True],

    'dense_units': [1, 1, 2, 2, 4, 4],
    'dense_activation': ['tanh', 'tanh', 'tanh', 'tanh', 'tanh', 'tanh',],
    'dense_trainable': [True, True, True, True, True, True],

    'LOSS': ['MSE', 'MAE', 'MSE', 'MAE', 'MSE', 'MAE'],
}

DNN_softmax_configs = {
    'dense0_units': [16, 16, 16, 16, 16],
    'dense0_activation': ['tanh', 'tanh', 'tanh', 'tanh', 'tanh'],
    'dense0_trainable': [True, True, True, True, True],
    'dense0_in': [True, True, True, True, True],

    'dense1_units': [8, 8, 8, 8, 8],
    'dense1_activation': ['elu', 'elu', 'elu', 'elu', 'elu'],
    'dense1_trainable': [True, True, True, True, True],
    'dense1_in': [True, True, True, True, True],

    'dense_units': [2, 2, 2, 2, 2],
    'dense_activation': ['softmax', 'softmax', 'softmax', 'softmax', 'softmax'],
    'dense_trainable': [True, True, True, True, True],

    'LOSS': ['MSE', 'MAE', 'MSLE', 'BCE', 'CCE'],
}


for configs in [P_sigmoid_configs, P_tanh_configs, P_softmax_configs, NN_sigmoid_configs, NN_tanh_configs, NN_softmax_configs, DNN_sigmoid_configs, DNN_tanh_configs, DNN_softmax_configs]:

    N_RUNS = len(configs['dense0_units'])

    for run in range(N_RUNS):
        args = []

        for k, v in configs.items():
            args.append(str(v[run]))
        args = ' '.join(args)

        os.system('python3 Dense_Dense.py %s' % args)
```
