## Supplementary material for "EpyNN: Educational python for Neural Networks": File S1: autotest_LSTM_Dense.py

```
"""Automatic testing.
"""
import os

### LSTM_Dense.py


lstm1_sigmoid_configs = {
    'lstm0_units': [4, 4, 4, 4, 4, 4, 4, 4, 4],
    'lstm0_activation': ['tanh', 'tanh', 'tanh', 'tanh', 'tanh', 'tanh', 'tanh', 'tanh', 'tanh'],
    'lstm0_sequences': [True, True, True, True, True, True, True, True, True],
    'lstm0_trainable': [True, True, True, True, True, True, True, True, True],
    'lstm0_in': [False, False, False, False, False, False, False, False, False],

    'lstm1_units': [2, 2, 2, 2, 2, 2, 2, 2, 2],
    'lstm1_activation': ['tanh', 'tanh', 'tanh', 'tanh', 'tanh', 'tanh', 'tanh', 'tanh', 'tanh'],
    'lstm1_sequences': [False, False, False, False, False, False, False, False, False],
    'lstm1_trainable': [True, True, True, True, True, True, True, True, True],
    'lstm1_in': [True, True, True, True, True, True, True, True, True],

    'dense_units': [1, 1, 1, 2, 2, 2, 4, 4, 4],
    'dense_activation': ['sigmoid', 'sigmoid', 'sigmoid', 'sigmoid', 'sigmoid', 'sigmoid', 'sigmoid', 'sigmoid', 'sigmoid'],
    'dense_trainable': [False, False, False, False, False, False, False, False, False],

    'LOSS': ['MSE', 'MAE', 'MSLE', 'MSE', 'MAE', 'MSLE', 'MSE', 'MAE', 'MSLE'],
}

lstm1_tanh_configs = {
    'lstm0_units': [4, 4, 4, 4, 4, 4],
    'lstm0_activation': ['tanh', 'tanh', 'tanh', 'tanh', 'tanh', 'tanh'],
    'lstm0_sequences': [True, True, True, True, True, True],
    'lstm0_trainable': [True, True, True, True, True, True],
    'lstm0_in': [False, False, False, False, False, False],

    'lstm1_units': [2, 2, 2, 2, 2, 2],
    'lstm1_activation': ['tanh', 'tanh', 'tanh', 'tanh', 'tanh', 'tanh'],
    'lstm1_sequences': [False, False, False, False, False, False],
    'lstm1_trainable': [True, True, True, True, True, True],
    'lstm1_in': [True, True, True, True, True, True],

    'dense_units': [1, 1, 2, 2, 4, 4],
    'dense_activation': ['tanh', 'tanh', 'tanh', 'tanh', 'tanh', 'tanh',],
    'dense_trainable': [False, False, False, False, False, False],

    'LOSS': ['MSE', 'MAE', 'MSE', 'MAE', 'MSE', 'MAE'],
}

lstm1_softmax_configs = {
    'lstm0_units': [4, 4, 4, 4, 4],
    'lstm0_activation': ['tanh', 'tanh', 'tanh', 'tanh', 'tanh'],
    'lstm0_sequences': [True, True, True, True, True],
    'lstm0_trainable': [True, True, True, True, True],
    'lstm0_in': [False, False, False, False, False],

    'lstm1_units': [2, 2, 2, 2, 2],
    'lstm1_activation': ['tanh', 'tanh', 'tanh', 'tanh', 'tanh'],
    'lstm1_sequences': [False, False, False, False, False],
    'lstm1_trainable': [True, True, True, True, True],
    'lstm1_in': [True, True, True, True, True],

    'dense_units': [2, 2, 2, 2, 2],
    'dense_activation': ['softmax', 'softmax', 'softmax', 'softmax', 'softmax'],
    'dense_trainable': [False, False, False, False, False],

    'LOSS': ['MSE', 'MAE', 'MSLE', 'BCE', 'CCE'],
}


lstm01_sigmoid_configs = {
    'lstm0_units': [4, 4, 4, 4, 4, 4, 4, 4, 4],
    'lstm0_activation': ['tanh', 'tanh', 'tanh', 'tanh', 'tanh', 'tanh', 'tanh', 'tanh', 'tanh'],
    'lstm0_sequences': [True, True, True, True, True, True, True, True, True],
    'lstm0_trainable': [True, True, True, True, True, True, True, True, True],
    'lstm0_in': [True, True, True, True, True, True, True, True, True],

    'lstm1_units': [2, 2, 2, 2, 2, 2, 2, 2, 2],
    'lstm1_activation': ['tanh', 'tanh', 'tanh', 'tanh', 'tanh', 'tanh', 'tanh', 'tanh', 'tanh'],
    'lstm1_sequences': [False, False, False, False, False, False, False, False, False],
    'lstm1_trainable': [True, True, True, True, True, True, True, True, True],
    'lstm1_in': [True, True, True, True, True, True, True, True, True],

    'dense_units': [1, 1, 1, 2, 2, 2, 4, 4, 4],
    'dense_activation': ['sigmoid', 'sigmoid', 'sigmoid', 'sigmoid', 'sigmoid', 'sigmoid', 'sigmoid', 'sigmoid', 'sigmoid'],
    'dense_trainable': [False, False, False, False, False, False, False, False, False],

    'LOSS': ['MSE', 'MAE', 'MSLE', 'MSE', 'MAE', 'MSLE', 'MSE', 'MAE', 'MSLE'],
}

lstm01_tanh_configs = {
    'lstm0_units': [4, 4, 4, 4, 4, 4],
    'lstm0_activation': ['tanh', 'tanh', 'tanh', 'tanh', 'tanh', 'tanh'],
    'lstm0_sequences': [True, True, True, True, True, True],
    'lstm0_trainable': [True, True, True, True, True, True],
    'lstm0_in': [True, True, True, True, True, True],

    'lstm1_units': [2, 2, 2, 2, 2, 2],
    'lstm1_activation': ['tanh', 'tanh', 'tanh', 'tanh', 'tanh', 'tanh'],
    'lstm1_sequences': [False, False, False, False, False, False],
    'lstm1_trainable': [True, True, True, True, True, True],
    'lstm1_in': [True, True, True, True, True, True],

    'dense_units': [1, 1, 2, 2, 4, 4],
    'dense_activation': ['tanh', 'tanh', 'tanh', 'tanh', 'tanh', 'tanh',],
    'dense_trainable': [False, False, False, False, False, False],

    'LOSS': ['MSE', 'MAE', 'MSE', 'MAE', 'MSE', 'MAE'],
}

lstm01_softmax_configs = {
    'lstm0_units': [4, 4, 4, 4, 4],
    'lstm0_activation': ['tanh', 'tanh', 'tanh', 'tanh', 'tanh'],
    'lstm0_sequences': [True, True, True, True, True],
    'lstm0_trainable': [True, True, True, True, True],
    'lstm0_in': [True, True, True, True, True],

    'lstm1_units': [2, 2, 2, 2, 2],
    'lstm1_activation': ['tanh', 'tanh', 'tanh', 'tanh', 'tanh'],
    'lstm1_sequences': [False, False, False, False, False],
    'lstm1_trainable': [True, True, True, True, True],
    'lstm1_in': [True, True, True, True, True],

    'dense_units': [2, 2, 2, 2, 2],
    'dense_activation': ['softmax', 'softmax', 'softmax', 'softmax', 'softmax'],
    'dense_trainable': [False, False, False, False, False],

    'LOSS': ['MSE', 'MAE', 'MSLE', 'BCE', 'CCE'],
}

sigmoid_configs = {
    'lstm0_units': [4, 4, 4, 4, 4, 4, 4, 4, 4],
    'lstm0_activation': ['tanh', 'tanh', 'tanh', 'tanh', 'tanh', 'tanh', 'tanh', 'tanh', 'tanh'],
    'lstm0_sequences': [True, True, True, True, True, True, True, True, True],
    'lstm0_trainable': [True, True, True, True, True, True, True, True, True],
    'lstm0_in': [True, True, True, True, True, True, True, True, True],

    'lstm1_units': [2, 2, 2, 2, 2, 2, 2, 2, 2],
    'lstm1_activation': ['tanh', 'tanh', 'tanh', 'tanh', 'tanh', 'tanh', 'tanh', 'tanh', 'tanh'],
    'lstm1_sequences': [True, True, True, True, True, True, True, True, True],
    'lstm1_trainable': [True, True, True, True, True, True, True, True, True],
    'lstm1_in': [True, True, True, True, True, True, True, True, True],

    'dense_units': [1, 1, 1, 2, 2, 2, 4, 4, 4],
    'dense_activation': ['sigmoid', 'sigmoid', 'sigmoid', 'sigmoid', 'sigmoid', 'sigmoid', 'sigmoid', 'sigmoid', 'sigmoid'],
    'dense_trainable': [False, False, False, False, False, False, False, False, False],

    'LOSS': ['MSE', 'MAE', 'MSLE', 'MSE', 'MAE', 'MSLE', 'MSE', 'MAE', 'MSLE'],
}

tanh_configs = {
    'lstm0_units': [4, 4, 4, 4, 4, 4],
    'lstm0_activation': ['tanh', 'tanh', 'tanh', 'tanh', 'tanh', 'tanh'],
    'lstm0_sequences': [True, True, True, True, True, True],
    'lstm0_trainable': [True, True, True, True, True, True],
    'lstm0_in': [True, True, True, True, True, True],

    'lstm1_units': [2, 2, 2, 2, 2, 2],
    'lstm1_activation': ['tanh', 'tanh', 'tanh', 'tanh', 'tanh', 'tanh'],
    'lstm1_sequences': [True, True, True, True, True, True],
    'lstm1_trainable': [True, True, True, True, True, True],
    'lstm1_in': [True, True, True, True, True, True],

    'dense_units': [1, 1, 2, 2, 4, 4],
    'dense_activation': ['tanh', 'tanh', 'tanh', 'tanh', 'tanh', 'tanh',],
    'dense_trainable': [False, False, False, False, False, False],

    'LOSS': ['MSE', 'MAE', 'MSE', 'MAE', 'MSE', 'MAE'],
}

softmax_configs = {
    'lstm0_units': [4, 4, 4, 4, 4],
    'lstm0_activation': ['tanh', 'tanh', 'tanh', 'tanh', 'tanh'],
    'lstm0_sequences': [True, True, True, True, True],
    'lstm0_trainable': [True, True, True, True, True],
    'lstm0_in': [True, True, True, True, True],

    'lstm1_units': [2, 2, 2, 2, 2],
    'lstm1_activation': ['tanh', 'tanh', 'tanh', 'tanh', 'tanh'],
    'lstm1_sequences': [True, True, True, True, True],
    'lstm1_trainable': [True, True, True, True, True],
    'lstm1_in': [True, True, True, True, True],

    'dense_units': [2, 2, 2, 2, 2],
    'dense_activation': ['softmax', 'softmax', 'softmax', 'softmax', 'softmax'],
    'dense_trainable': [False, False, False, False, False],

    'LOSS': ['MSE', 'MAE', 'MSLE', 'BCE', 'CCE'],
}


for configs in [lstm1_sigmoid_configs, lstm1_tanh_configs, lstm1_softmax_configs, lstm01_sigmoid_configs, lstm01_tanh_configs, lstm01_softmax_configs, sigmoid_configs, tanh_configs, softmax_configs]:

    N_RUNS = len(configs['lstm0_units'])

    for run in range(N_RUNS):
        args = []

        for k, v in configs.items():
            args.append(str(v[run]))
        args = ' '.join(args)

        os.system('python3 LSTM_Dense.py %s' % args)
```
