## Supplementary material for "EpyNN: Educational python for Neural Networks": File S1: Conv_Pool_Dense.py

```
"""
Config is strictly identical between Keras and EpyNN.
Parameters in ### SHARED CONSTANTS ### are tunable and propagate to Keras and EpyNN.
Weights/bias are initialized with Keras and used for Keras and EpyNN.
See in ./utils.py for more code.

Architecture is:
    Conv > Pool > Conv > Pool > Dense
"""
####################### SHARED CONSTANTS ######################
# Data
N_SAMPLES = 100
HEIGHT = 128
WIDTH = 128
DEPTH = 4

BATCH_SIZE = N_SAMPLES # We don't make any batch here

# Layers
conv0_filters = 3
conv0_kernel = (2, 4)
conv0_strides = (4, 2)
conv0_activation = 'relu'
conv0_bias = True
conv0_trainable = True
conv0_in = True

pool0_kernel = (2, 4)
pool0_strides = (2, 4)
pool0_in = True

conv1_filters = 6
conv1_kernel = (4, 2)
conv1_strides = (4, 2)
conv1_activation = 'relu'
conv1_bias = True
conv1_trainable = True
conv1_in = True

pool1_kernel = (2, 2)
pool1_strides = (2, 2)
pool1_in = True

dense_units = 2
dense_activation = 'sigmoid'
dense_trainable = False

# Run settings
N_EPOCHS = 2
LRATE = 0.01

LOSS = 'MSE'

# Support for autotest_Conv_Pool_Dense.py
import sys

if len(sys.argv) > 1:
    del sys.argv[0]
    args = sys.argv
    args = [int(arg) if arg.isdigit() else arg for arg in args]
    args = [True if arg == 'True' else arg for arg in args]
    args = [False if arg == 'False' else arg for arg in args]
    conv0_filters, conv0_in, pool0_in, conv1_filters, conv1_in, pool1_in, dense_units, dense_activation, dense_trainable, LOSS = args

######################## SEEDING/CFG ##########################
from timeit import default_timer as timer
from termcolor import cprint
import numpy as np
import random
import os

# CPU run - Disable GPU
os.environ["CUDA_DEVICE_ORDER"] = "PCI_BUS_ID"
os.environ["CUDA_VISIBLE_DEVICES"] = ""
# Hide tensorflow debug info on call
os.environ['TF_CPP_MIN_LOG_LEVEL'] = '3'

# Seeding everywhere
import tensorflow as tf

SEED = 1
random.seed(SEED)
np.random.seed(SEED)
tf.random.set_seed(SEED)

# Float precision for TF (NumPy defaults to float64)
tf.keras.backend.set_floatx('float64')

# NumPy behavior on float arithmetics errors
np.seterr(all='raise')

######################## SHARED DATA ##########################
from epynn.commons.io import scale_features
from utils import log_data

X_train = np.random.uniform(0, 1, (N_SAMPLES, HEIGHT, WIDTH, DEPTH))
Y_train = np.random.randint(0, dense_units, (N_SAMPLES))

# One-hot encoding
Y_train = np.eye(dense_units)[list(Y_train)]

###############################################################
##################### TENSORFLOW/KERAS ########################
from tensorflow.keras import Sequential
from tensorflow.keras.layers import Conv2D as KConv
from tensorflow.keras.layers import MaxPooling2D as KPool
from tensorflow.keras.layers import Flatten as KFlatten
from tensorflow.keras.layers import Dense as KDense
from utils import get_losses

keras_loss = get_losses(LOSS)
keras_opt = tf.keras.optimizers.SGD(learning_rate=LRATE)

# Layers
keras_conv0 = KConv(conv0_filters, kernel_size=conv0_kernel, strides=conv0_strides, activation=conv0_activation, use_bias=conv0_bias)
keras_pool0 = KPool(pool_size=pool0_kernel, strides=pool0_strides)
keras_conv1 = KConv(conv1_filters, kernel_size=conv1_kernel, strides=conv1_strides, activation=conv1_activation, use_bias=conv1_bias)
keras_pool1 = KPool(pool_size=pool1_kernel, strides=pool1_strides)

keras_flatten = KFlatten()
keras_dense = KDense(dense_units, activation=dense_activation)

# Model
keras_model = Sequential()
if conv0_in:
    keras_model.add(keras_conv0)
    keras_conv0.trainable = False
if pool0_in:
    keras_model.add(keras_pool0)
    keras_pool0.trainable = False
if conv1_in:
    keras_model.add(keras_conv1)
    keras_conv1.trainable = False
if pool1_in:
    keras_model.add(keras_pool1)
    keras_pool1.trainable = False

keras_model.add(keras_flatten)
keras_model.add(keras_dense)
keras_dense.trainable = False

# Init
keras_model.compile(optimizer=keras_opt, loss=keras_loss, metrics=['accuracy'])
keras_model.fit(X_train, Y_train, epochs=N_EPOCHS, batch_size=BATCH_SIZE, verbose=0)

################# SHARED PARAMETERS
if conv0_in:
    conv0_W = keras_conv0.get_weights()[0]
    conv0_b = keras_conv0.get_weights()[1] if conv0_bias else np.zeros(conv0_filters)

if conv1_in:
    conv1_W = keras_conv1.get_weights()[0]
    conv1_b = keras_conv1.get_weights()[1] if conv1_bias else np.zeros(conv1_filters)

dense_W, dense_b = keras_dense.get_weights()

###################### RESUME
###################### TENSORFLOW/KERAS TRAINING
keras_conv0.trainable = conv0_trainable
keras_conv1.trainable = conv1_trainable
keras_dense.trainable = dense_trainable

keras_start = timer()
keras_model.compile(optimizer=keras_opt, loss=keras_loss, metrics=['accuracy'])
keras_model.fit(X_train, Y_train, epochs=N_EPOCHS, batch_size=BATCH_SIZE, verbose=1)
keras_end = timer() - keras_start
#############################

keras_model.summary()

keras_preds = keras_model.predict(X_train)

###############################################################
############################# EpyNN ###########################
from epynn.dense.models import Dense as EDense
from epynn.embedding.models import Embedding
from epynn.network.models import EpyNN
from epynn.convolution.models import Convolution as EConv
from epynn.pooling.models import Pooling as EPool
from epynn.flatten.models import Flatten as EFlatten
from utils import get_activation
from settings import se_hPars

se_hPars['learning_rate'] = LRATE
conv0_activation = get_activation(conv0_activation)
conv1_activation = get_activation(conv1_activation)
dense_activation = get_activation(dense_activation)

# Layers
epynn_embedding = Embedding(X_train, Y_train, relative_size=(1, 0, 0), batch_size=BATCH_SIZE)

epynn_conv0 = EConv(conv0_filters, filter_size=conv0_kernel, strides=conv0_strides, activate=conv0_activation, use_bias=conv0_bias)
epynn_pool0 = EPool(pool_size=pool0_kernel, strides=pool0_strides)
epynn_conv1 = EConv(conv1_filters, filter_size=conv1_kernel, strides=conv1_strides, activate=conv1_activation, use_bias=conv1_bias)
epynn_pool1 = EPool(pool_size=pool1_kernel, strides=pool1_strides)

epynn_flatten = EFlatten()
epynn_dense = EDense(units=dense_units, activate=dense_activation)

# Model
layers = [epynn_embedding]

if conv0_in:
    epynn_conv0.p['W'] = conv0_W
    epynn_conv0.p['b'] = conv0_b
    epynn_conv0.trainable = conv0_trainable
    layers.append(epynn_conv0)

if pool0_in:
    layers.append(epynn_pool0)

if conv1_in:
    epynn_conv1.p['W'] = conv1_W
    epynn_conv1.p['b'] = conv1_b
    epynn_conv1.trainable = conv1_trainable
    layers.append(epynn_conv1)

if pool1_in:
    layers.append(epynn_pool1)

layers.append(epynn_flatten)

epynn_dense.p['W'] = dense_W
epynn_dense.p['b'] = dense_b
epynn_dense.trainable = dense_trainable
layers.append(epynn_dense)

# Init
epynn_model = EpyNN(layers=layers)
epynn_model.initialize(loss=LOSS, se_hPars=se_hPars.copy(), seed=SEED, params=False, end='\n')

### EpyNN TRAINING ###
epynn_start = timer()
epynn_model.train(epochs=N_EPOCHS, verbose=1, init_logs=False)
epynn_end = timer() - epynn_start
######################

epynn_preds = epynn_model.forward(X_train)


###############################################################
############################ Compare ##########################
from utils import print_logs

data = (Y_train, epynn_preds, keras_preds, LOSS, keras_loss, epynn_model, keras_end, epynn_end)

kloss_RMSD, preds_RMSD = print_logs(data, extended=False)

if len(sys.argv) > 1:
    args = [str(arg) for arg in args]
    msg = str(N_SAMPLES)+', '+str(HEIGHT)+', '+str(WIDTH)+', '+str(DEPTH)+', '+', '.join(args)+', '+str(kloss_RMSD)+', '+str(preds_RMSD)
    with open('autotest_Conv_Pool_Dense.dat', 'a') as f:
        f.write(msg)
        f.write('\n')
```
