## Supplementary material for "EpyNN: Educational python for Neural Networks": File S1: Dense_Dense.py

```
"""
Config is strictly identical between Keras and EpyNN.
Parameters in ### SHARED CONSTANTS ### are tunable and propagate to Keras and EpyNN.
Weights/bias are initialized with Keras and used for Keras and EpyNN.
See in ./utils.py for more code.

Architecture is:
    Dense > Dense > Dense
"""
####################### SHARED CONSTANTS ######################
# Data
N_SAMPLES = 32
N_FEATURES = 64


BATCH_SIZE = N_SAMPLES # We don't make any batch here

# Layers
dense0_units = 16
dense0_activation = 'tanh'
dense0_trainable = True
dense0_in = True

dense1_units = 8
dense1_activation = 'elu'
dense1_trainable = True
dense1_in = True

dense_units = 4
dense_activation = 'softmax'
dense_trainable = True

# Run settings
N_EPOCHS = 2
LRATE = 0.01

LOSS = 'MSE'

# Support for autotest_Dense_Dense.py
import sys

if len(sys.argv) > 1:
    del sys.argv[0]
    args = sys.argv
    args = [int(arg) if arg.isdigit() else arg for arg in args]
    args = [True if arg == 'True' else arg for arg in args]
    args = [False if arg == 'False' else arg for arg in args]
    dense0_units, dense0_activation, dense0_trainable, dense0_in, dense1_units, dense1_activation, dense1_trainable, dense1_in, dense_units, dense_activation, dense_trainable, LOSS = args

######################## SEEDING/CFG ##########################
from timeit import default_timer as timer
from termcolor import cprint
import numpy as np
import random
import os

# CPU run - Disable GPU
os.environ["CUDA_DEVICE_ORDER"] = "PCI_BUS_ID"
os.environ["CUDA_VISIBLE_DEVICES"] = ""
# Hide tensorflow debug info on call
os.environ['TF_CPP_MIN_LOG_LEVEL'] = '3'

# Seeding everywhere
import tensorflow as tf

SEED = 1
random.seed(SEED)
np.random.seed(SEED)
tf.random.set_seed(SEED)

# Float precision for TF (NumPy defaults to float64)
tf.keras.backend.set_floatx('float64')

# NumPy behavior on float arithmetics errors
np.seterr(all='raise')

######################## SHARED DATA ##########################
from utils import log_data

# Prepare dummy data
X_train = np.random.randint(-2, 2, (N_SAMPLES, N_FEATURES)).astype('float64')
Y_train = np.random.randint(0, dense_units, (N_SAMPLES))

# One-hot encode X_train and Y_train
Y_train = np.eye(dense_units)[Y_train]

# Logs
log_data(X_train, Y_train)

###############################################################
##################### TENSORFLOW/KERAS ########################
from tensorflow.keras import Sequential
from tensorflow.keras.layers import Dense as KDense
from utils import get_losses


keras_loss = get_losses(LOSS)
keras_opt = tf.keras.optimizers.SGD(learning_rate=LRATE)

# Layers
keras_dense0 = KDense(dense0_units, activation=dense0_activation)
keras_dense1 = KDense(dense1_units, activation=dense1_activation)

keras_dense = KDense(dense_units, activation=dense_activation)

# Model
keras_model = Sequential()
if dense0_in:
    keras_model.add(keras_dense0)
    keras_dense0.trainable = False
if dense1_in:
    keras_model.add(keras_dense1)
    keras_dense1.trainable = False

keras_model.add(keras_dense)
keras_dense.trainable = False

# Init
keras_model.compile(optimizer=keras_opt, loss=keras_loss, metrics=['accuracy'])
keras_model.fit(X_train, Y_train, epochs=N_EPOCHS, batch_size=BATCH_SIZE, verbose=1)

################# SHARED PARAMETERS

if dense0_in:
    dense0_W, dense0_b = keras_dense0.get_weights()

if dense1_in:
    dense1_W, dense1_b = keras_dense1.get_weights()

dense_W, dense_b = keras_dense.get_weights()

###################### RESUME
###################### TENSORFLOW/KERAS TRAINING
keras_dense0.trainable = dense0_trainable
keras_dense1.trainable = dense1_trainable
keras_dense.trainable = dense_trainable

keras_start = timer()
keras_model.compile(optimizer=keras_opt, loss=keras_loss, metrics=['accuracy'])
keras_model.fit(X_train, Y_train, epochs=N_EPOCHS, batch_size=BATCH_SIZE, verbose=1)
keras_end = timer() - keras_start
#############################
keras_model.summary()

keras_preds = keras_model.predict(X_train)

###############################################################
############################# EpyNN ###########################
from epynn.commons.maths import tanh
from epynn.dense.models import Dense as EDense
from epynn.embedding.models import Embedding
from epynn.network.models import EpyNN
from utils import get_activation
from settings import se_hPars

se_hPars['learning_rate'] = LRATE
dense0_activation = get_activation(dense0_activation)
dense1_activation = get_activation(dense1_activation)
dense_activation = get_activation(dense_activation)

# Layers
epynn_embedding = Embedding(X_train, Y_train, relative_size=(1, 0, 0), batch_size=BATCH_SIZE)
epynn_dense0 = EDense(units=dense0_units, activate=dense0_activation)
epynn_dense1 = EDense(units=dense1_units, activate=dense1_activation)

epynn_dense = EDense(units=dense_units, activate=dense_activation)

# Model
layers = [epynn_embedding]

if dense0_in:
    epynn_dense0.p['W'] = dense0_W
    epynn_dense0.p['b'] = dense0_b
    epynn_dense0.trainable = dense0_trainable
    layers.append(epynn_dense0)

if dense1_in:
    epynn_dense1.p['W'] = dense1_W
    epynn_dense1.p['b'] = dense1_b
    epynn_dense1.trainable = dense1_trainable
    layers.append(epynn_dense1)

epynn_dense.p['W'] = dense_W
epynn_dense.p['b'] = dense_b
epynn_dense.trainable = dense_trainable
layers.append(epynn_dense)

# Init
epynn_model = EpyNN(layers=layers)
epynn_model.initialize(loss=LOSS, se_hPars=se_hPars.copy(), seed=SEED, params=False, end='\r')

### EpyNN TRAINING ###
epynn_start = timer()
epynn_model.train(epochs=N_EPOCHS, verbose=1, init_logs=False)
epynn_end = timer() - epynn_start
######################

epynn_preds = epynn_model.forward(X_train)


###############################################################
############################ Compare ##########################
from utils import print_logs

data = (Y_train, epynn_preds, keras_preds, LOSS, keras_loss, epynn_model, keras_end, epynn_end)

kloss_RMSD, preds_RMSD = print_logs(data)

if len(sys.argv) > 1:
    args = [str(arg) for arg in args]
    msg = str(N_SAMPLES)+', '+str(N_FEATURES)+', '+', '.join(args)+', '+str(kloss_RMSD)+', '+str(preds_RMSD)
    with open('autotest_Dense_Dense.dat', 'a') as f:
        f.write(msg)
        f.write('\n')
```
