## Supplementary material for "EpyNN: Educational python for Neural Networks": File S1: LSTM_Dense.py

```
"""
Config is strictly identical between Keras and EpyNN.
Parameters in ### SHARED CONSTANTS ### are tunable and propagate to Keras and EpyNN.
Weights/bias are initialized with Keras and used for Keras and EpyNN.
See in ./utils.py for more code.

Architecture is:
    LSTM > LSTM > Dense
"""
####################### SHARED CONSTANTS ######################
# Data
N_SAMPLES = 32
STEPS = 10
VSIZE = 5

BATCH_SIZE = N_SAMPLES # We don't make any batch here

# Layers
lstm0_units = 4
lstm0_activation = 'tanh'
lstm0_sequences = True
lstm0_trainable = True
lstm0_in = True

lstm1_units = 2
lstm1_activation = 'tanh'
lstm1_sequences = False
lstm1_trainable = True
lstm1_in = True

dense_units = 2
dense_activation = 'softmax'
dense_trainable = False

# Run settings
N_EPOCHS = 2
LRATE = 0.01

LOSS = 'CCE'

# Support for autotest_LSTM_Dense.py
import sys

if len(sys.argv) > 1:
    del sys.argv[0]
    args = sys.argv
    args = [int(arg) if arg.isdigit() else arg for arg in args]
    args = [True if arg == 'True' else arg for arg in args]
    args = [False if arg == 'False' else arg for arg in args]
    lstm0_units, lstm0_activation, lstm0_sequences, lstm0_trainable, lstm0_in, lstm1_units, lstm1_activation, lstm1_sequences, lstm1_trainable, lstm1_in, dense_units, dense_activation, dense_trainable, LOSS = args

######################## SEEDING/CFG ##########################
from timeit import default_timer as timer
from termcolor import cprint
import numpy as np
import random
import os

# CPU run - Disable GPU
os.environ["CUDA_DEVICE_ORDER"] = "PCI_BUS_ID"
os.environ["CUDA_VISIBLE_DEVICES"] = ""
# Hide tensorflow debug info on call
os.environ['TF_CPP_MIN_LOG_LEVEL'] = '3'

# Seeding everywhere
import tensorflow as tf

SEED = 1
random.seed(SEED)
np.random.seed(SEED)
tf.random.set_seed(SEED)

# Float precision for TF (NumPy defaults to float64)
tf.keras.backend.set_floatx('float64')

# NumPy behavior on float arithmetics errors
np.seterr(all='raise')

######################## SHARED DATA ##########################
from utils import log_data

# Prepare dummy data
X_train = np.random.randint(-2, 2, (N_SAMPLES, STEPS, VSIZE)).astype('float64')
Y_train = np.random.randint(0, dense_units, (N_SAMPLES))

# One-hot encode X_train and Y_train
Y_train = np.eye(dense_units)[Y_train]

# Logs
log_data(X_train, Y_train)

###############################################################
##################### TENSORFLOW/KERAS ########################
from tensorflow.keras import Sequential
from tensorflow.keras.layers import Dense as KDense
from tensorflow.keras.layers import LSTM as KLSTM
from tensorflow.keras.layers import Flatten as KFlatten
from utils import get_losses


keras_loss = get_losses(LOSS)
keras_opt = tf.keras.optimizers.SGD(learning_rate=LRATE)

# Layers
keras_lstm0 = KLSTM(lstm0_units, activation=lstm0_activation, return_sequences=lstm0_sequences, recurrent_activation='sigmoid')
keras_lstm1 = KLSTM(lstm1_units, activation=lstm1_activation, return_sequences=lstm1_sequences, recurrent_activation='sigmoid')

keras_flatten = KFlatten()
keras_dense = KDense(dense_units, activation=dense_activation)

# Model
keras_model = Sequential()
if lstm0_in:
    keras_model.add(keras_lstm0)
    keras_lstm0.trainable = False
if lstm1_in:
    keras_model.add(keras_lstm1)
    keras_lstm1.trainable = False

keras_model.add(keras_flatten)

keras_dense.trainable = dense_trainable
keras_model.add(keras_dense)

# Init
keras_model.compile(optimizer=keras_opt, loss=keras_loss, metrics=['accuracy'])
keras_model.fit(X_train, Y_train, epochs=N_EPOCHS, batch_size=BATCH_SIZE, verbose=1)

################# SHARED PARAMETERS

if lstm0_in:
    lstm0_Ui, lstm0_Uf, lstm0_Ug, lstm0_Uo = np.split(keras_lstm0.get_weights()[0], 4, axis=1)
    lstm0_Vi, lstm0_Vf, lstm0_Vg, lstm0_Vo = np.split(keras_lstm0.get_weights()[1], 4, axis=1)
    lstm0_bi, lstm0_bf, lstm0_bg, lstm0_bo = np.split(keras_lstm0.get_weights()[2], 4, axis=0)

if lstm1_in:
    lstm1_Ui, lstm1_Uf, lstm1_Ug, lstm1_Uo = np.split(keras_lstm1.get_weights()[0], 4, axis=1)
    lstm1_Vi, lstm1_Vf, lstm1_Vg, lstm1_Vo = np.split(keras_lstm1.get_weights()[1], 4, axis=1)
    lstm1_bi, lstm1_bf, lstm1_bg, lstm1_bo = np.split(keras_lstm1.get_weights()[2], 4, axis=0)

dense_W, dense_b = keras_dense.get_weights()
###################### RESUME
###################### TENSORFLOW/KERAS TRAINING
keras_lstm0.trainable = lstm0_trainable
keras_lstm1.trainable = lstm1_trainable
keras_dense.trainable = dense_trainable

keras_start = timer()
keras_model.compile(optimizer=keras_opt, loss=keras_loss, metrics=['accuracy'])
keras_model.fit(X_train, Y_train, epochs=N_EPOCHS, batch_size=BATCH_SIZE, verbose=1)
keras_end = timer() - keras_start
#############################
keras_model.summary()

keras_preds = keras_model.predict(X_train)

###############################################################
############################# EpyNN ###########################
from epynn.commons.maths import tanh
from epynn.dense.models import Dense as EDense
from epynn.lstm.models import LSTM as ELSTM
from epynn.embedding.models import Embedding
from epynn.network.models import EpyNN
from epynn.flatten.models import Flatten as EFlatten
from utils import get_activation
from settings import se_hPars

se_hPars['learning_rate'] = LRATE
lstm0_activation = get_activation(lstm0_activation)
lstm1_activation = get_activation(lstm1_activation)
dense_activation = get_activation(dense_activation)

# Layers
epynn_embedding = Embedding(X_train, Y_train, relative_size=(1, 0, 0), batch_size=BATCH_SIZE)
epynn_lstm0 = ELSTM(unit_cells=lstm0_units, sequences=lstm0_sequences, activate=lstm0_activation, clip_gradients=False)
epynn_lstm1 = ELSTM(unit_cells=lstm1_units, sequences=lstm1_sequences, activate=lstm1_activation, clip_gradients=False)

epynn_flatten = EFlatten()
epynn_dense = EDense(units=dense_units, activate=dense_activation)

# Model
layers = [epynn_embedding]

if lstm0_in:
    epynn_lstm0.p['Ui'] = lstm0_Ui
    epynn_lstm0.p['Vi'] = lstm0_Vi
    epynn_lstm0.p['bi'] = lstm0_bi
    epynn_lstm0.p['Uf'] = lstm0_Uf
    epynn_lstm0.p['Vf'] = lstm0_Vf
    epynn_lstm0.p['bf'] = lstm0_bf
    epynn_lstm0.p['Ug'] = lstm0_Ug
    epynn_lstm0.p['Vg'] = lstm0_Vg
    epynn_lstm0.p['bg'] = lstm0_bg
    epynn_lstm0.p['Uo'] = lstm0_Uo
    epynn_lstm0.p['Vo'] = lstm0_Vo
    epynn_lstm0.p['bo'] = lstm0_bo
    epynn_lstm0.trainable = lstm0_trainable
    layers.append(epynn_lstm0)

if lstm1_in:
    epynn_lstm1.p['Ui'] = lstm1_Ui
    epynn_lstm1.p['Vi'] = lstm1_Vi
    epynn_lstm1.p['bi'] = lstm1_bi
    epynn_lstm1.p['Uf'] = lstm1_Uf
    epynn_lstm1.p['Vf'] = lstm1_Vf
    epynn_lstm1.p['bf'] = lstm1_bf
    epynn_lstm1.p['Ug'] = lstm1_Ug
    epynn_lstm1.p['Vg'] = lstm1_Vg
    epynn_lstm1.p['bg'] = lstm1_bg
    epynn_lstm1.p['Uo'] = lstm1_Uo
    epynn_lstm1.p['Vo'] = lstm1_Vo
    epynn_lstm1.p['bo'] = lstm1_bo
    epynn_lstm1.trainable = lstm1_trainable
    layers.append(epynn_lstm1)

layers.append(epynn_flatten)

epynn_dense.p['W'] = dense_W
epynn_dense.p['b'] = dense_b
epynn_dense.trainable = dense_trainable
layers.append(epynn_dense)

# Init
epynn_model = EpyNN(layers=layers)
epynn_model.initialize(loss=LOSS, se_hPars=se_hPars.copy(), seed=SEED, params=False, end='\r')

### EpyNN TRAINING ###
epynn_start = timer()
epynn_model.train(epochs=N_EPOCHS, verbose=1, init_logs=False)
epynn_end = timer() - epynn_start
######################

epynn_preds = epynn_model.forward(X_train)


###############################################################
############################ Compare ##########################
from utils import print_logs

data = (Y_train, epynn_preds, keras_preds, LOSS, keras_loss, epynn_model, keras_end, epynn_end)

kloss_RMSD, preds_RMSD = print_logs(data)

if len(sys.argv) > 1:
    args = [str(arg) for arg in args]
    msg = str(N_SAMPLES)+', '+str(STEPS)+', '+str(VSIZE)+', '+', '.join(args)+', '+str(kloss_RMSD)+', '+str(preds_RMSD)
    with open('autotest_LSTM_Dense.dat', 'a') as f:
        f.write(msg)
        f.write('\n')
```
