## Supplementary material for "EpyNN: Educational python for Neural Networks": File S1: README

Contents:
    1. Dense_Dense.py: Code for Feed-Forward network comparison EpyNN vs TF/Keras
    2. RNN_Dense.py: Idem with RNN
    3. LSTM_Dense.py: Idem with LSTM
    4. GRU_Dense.py: Idem with GRU
    5. Conv_Pool_Dense.py: Idem with ConvPool blocks
    - utils.py: some functions used across 1-5.
    - Others:
        - autotest*py: One file for each 1-5. Perform batch tests with multiple configurations.
        - autotest*dat: Output of autotest*py.
        - run.sh: Loop over autotest*py and run them.
        - analysis: codes to process and plot autotest*dat.
            ./proc.sh
            python3 plot.py

Usage:
    python3 file.py
    bash file.sh

Note that:
    E_SAFE was set to 0 in EpyNN/epynn/commons/loss.py for the purpose of those experiments.

IMPORTANT:
    If you encounter the following issue with Tensorflow https://github.com/tensorflow/tensorflow/issues/47242 then downgrade NumPy to 1.19.4.
