## Supplementary material for "EpyNN: Educational python for Neural Networks": File S1: RNN_Dense.py

```
"""
Config is strictly identical between Keras and EpyNN.
Parameters in ### SHARED CONSTANTS ### are tunable and propagate to Keras and EpyNN.
Weights/bias are initialized with Keras and used for Keras and EpyNN.
See in ./utils.py for more code.

Architecture is:
    RNN > RNN > Dense
"""
####################### SHARED CONSTANTS ######################
# Data
N_SAMPLES = 32
STEPS = 10
VSIZE = 5

BATCH_SIZE = N_SAMPLES # We don't make any batch here

# Layers
rnn0_units = 4
rnn0_activation = 'tanh'
rnn0_sequences = True
rnn0_trainable = True
rnn0_in = True

rnn1_units = 2
rnn1_activation = 'tanh'
rnn1_sequences = False
rnn1_trainable = True
rnn1_in = True

dense_units = 2
dense_activation = 'sigmoid'
dense_trainable = False

# Run settings
N_EPOCHS = 2
LRATE = 0.01

LOSS = 'MSE'

# Support for autotest_RNN_Dense.py
import sys

if len(sys.argv) > 1:
    del sys.argv[0]
    args = sys.argv
    args = [int(arg) if arg.isdigit() else arg for arg in args]
    args = [True if arg == 'True' else arg for arg in args]
    args = [False if arg == 'False' else arg for arg in args]
    rnn0_units, rnn0_activation, rnn0_sequences, rnn0_trainable, rnn0_in, rnn1_units, rnn1_activation, rnn1_sequences, rnn1_trainable, rnn1_in, dense_units, dense_activation, dense_trainable, LOSS = args

######################## SEEDING/CFG ##########################
from timeit import default_timer as timer
from termcolor import cprint
import numpy as np
import random
import os

# CPU run - Disable GPU
os.environ["CUDA_DEVICE_ORDER"] = "PCI_BUS_ID"
os.environ["CUDA_VISIBLE_DEVICES"] = ""
# Hide tensorflow debug info on call
os.environ['TF_CPP_MIN_LOG_LEVEL'] = '3'

# Seeding everywhere
import tensorflow as tf

SEED = 1
random.seed(SEED)
np.random.seed(SEED)
tf.random.set_seed(SEED)

# Float precision for TF (NumPy defaults to float64)
tf.keras.backend.set_floatx('float64')

# NumPy behavior on float arithmetics errors
np.seterr(all='raise')

######################## SHARED DATA ##########################
from utils import log_data

# Prepare dummy data
X_train = np.random.randint(-2, 2, (N_SAMPLES, STEPS, VSIZE)).astype('float64')
Y_train = np.random.randint(0, dense_units, (N_SAMPLES))

# One-hot encode X_train and Y_train
Y_train = np.eye(dense_units)[Y_train]

# Logs
log_data(X_train, Y_train)

###############################################################
##################### TENSORFLOW/KERAS ########################
from tensorflow.keras import Sequential
from tensorflow.keras.layers import Dense as KDense
from tensorflow.keras.layers import SimpleRNN as KRNN
from tensorflow.keras.layers import Flatten as KFlatten
from utils import get_losses


keras_loss = get_losses(LOSS)
keras_opt = tf.keras.optimizers.SGD(learning_rate=LRATE)

# Layers
keras_rnn0 = KRNN(rnn0_units, activation=rnn0_activation, return_sequences=rnn0_sequences)
keras_rnn1 = KRNN(rnn1_units, activation=rnn1_activation, return_sequences=rnn1_sequences)

keras_flatten = KFlatten()
keras_dense = KDense(dense_units, activation=dense_activation)

# Model
keras_model = Sequential()
if rnn0_in:
    keras_model.add(keras_rnn0)
    keras_rnn0.trainable = False
if rnn1_in:
    keras_model.add(keras_rnn1)
    keras_rnn1.trainable = False

keras_model.add(keras_flatten)

keras_dense.trainable = dense_trainable
keras_model.add(keras_dense)

# Init
keras_model.compile(optimizer=keras_opt, loss=keras_loss, metrics=['accuracy'])
keras_model.fit(X_train, Y_train, epochs=N_EPOCHS, batch_size=BATCH_SIZE, verbose=1)

################# SHARED PARAMETERS

if rnn0_in:
    rnn0_U, rnn0_V, rnn0_b = keras_rnn0.get_weights()

if rnn1_in:
    rnn1_U, rnn1_V, rnn1_b = keras_rnn1.get_weights()

dense_W, dense_b = keras_dense.get_weights()
###################### RESUME
###################### TENSORFLOW/KERAS TRAINING
keras_rnn0.trainable = rnn0_trainable
keras_rnn1.trainable = rnn1_trainable
keras_dense.trainable = dense_trainable

keras_start = timer()
keras_model.compile(optimizer=keras_opt, loss=keras_loss, metrics=['accuracy'])
keras_model.fit(X_train, Y_train, epochs=N_EPOCHS, batch_size=BATCH_SIZE, verbose=1)
keras_end = timer() - keras_start
#############################
keras_model.summary()

keras_preds = keras_model.predict(X_train)

###############################################################
############################# EpyNN ###########################
from epynn.commons.maths import tanh
from epynn.dense.models import Dense as EDense
from epynn.rnn.models import RNN as ERNN
from epynn.embedding.models import Embedding
from epynn.network.models import EpyNN
from epynn.flatten.models import Flatten as EFlatten
from utils import get_activation
from settings import se_hPars

se_hPars['learning_rate'] = LRATE
rnn0_activation = get_activation(rnn0_activation)
rnn1_activation = get_activation(rnn1_activation)
dense_activation = get_activation(dense_activation)

# Layers
epynn_embedding = Embedding(X_train, Y_train, relative_size=(1, 0, 0), batch_size=BATCH_SIZE)
epynn_rnn0 = ERNN(unit_cells=rnn0_units, sequences=rnn0_sequences, activate=rnn0_activation, clip_gradients=False)
epynn_rnn1 = ERNN(unit_cells=rnn1_units, sequences=rnn1_sequences, activate=rnn1_activation, clip_gradients=False)

epynn_flatten = EFlatten()
epynn_dense = EDense(units=dense_units, activate=dense_activation)

# Model
layers = [epynn_embedding]

if rnn0_in:
    epynn_rnn0.p['U'] = rnn0_U
    epynn_rnn0.p['V'] = rnn0_V
    epynn_rnn0.p['b'] = rnn0_b
    epynn_rnn0.trainable = rnn0_trainable
    layers.append(epynn_rnn0)

if rnn1_in:
    epynn_rnn1.p['U'] = rnn1_U
    epynn_rnn1.p['V'] = rnn1_V
    epynn_rnn1.p['b'] = rnn1_b
    epynn_rnn1.trainable = rnn1_trainable
    layers.append(epynn_rnn1)

layers.append(epynn_flatten)

epynn_dense.p['W'] = dense_W
epynn_dense.p['b'] = dense_b
epynn_dense.trainable = dense_trainable
layers.append(epynn_dense)

# Init
epynn_model = EpyNN(layers=layers)
epynn_model.initialize(loss=LOSS, se_hPars=se_hPars.copy(), seed=SEED, params=False, end='\r')

### EpyNN TRAINING ###
epynn_start = timer()
epynn_model.train(epochs=N_EPOCHS, verbose=1, init_logs=False)
epynn_end = timer() - epynn_start
######################

epynn_preds = epynn_model.forward(X_train)


###############################################################
############################ Compare ##########################
from utils import print_logs

data = (Y_train, epynn_preds, keras_preds, LOSS, keras_loss, epynn_model, keras_end, epynn_end)

kloss_RMSD, preds_RMSD = print_logs(data)

if len(sys.argv) > 1:
    args = [str(arg) for arg in args]
    msg = str(N_SAMPLES)+', '+str(STEPS)+', '+str(VSIZE)+', '+', '.join(args)+', '+str(kloss_RMSD)+', '+str(preds_RMSD)
    with open('autotest_RNN_Dense.dat', 'a') as f:
        f.write(msg)
        f.write('\n')
```
