## Supplementary material for "EpyNN: Educational python for Neural Networks": File S1: settings.py

```
# EpyNN/epynn/settings.py


# HYPERPARAMETERS SETTINGS
se_hPars = {
    # Schedule learning rate
    'learning_rate': 0.01,
    'schedule': 'steady',
    'decay_k': 0,
    'cycle_epochs': 0,
    'cycle_descent': 0,
    # Tune activation function
    'ELU_alpha': 1,
    'LRELU_alpha': 0.3,
    'softmax_temperature': 1,
}
"""Hyperparameters dictionary settings.

Set hyperparameters for model and layer.
"""
```
