## Supplementary material for "EpyNN: Educational python for Neural Networks": File S1: utils.py

```
from epynn.commons.maths import sigmoid, tanh, softmax, relu, elu, identity, lrelu
import tensorflow as tf
from termcolor import cprint
import numpy as np

def log_data(X_train, Y_train):

    print('X_train.shape:', X_train.shape)
    print('X_train[0]:', X_train[0])
    print('Y_train.shape:',Y_train.shape)
    print('Y_train[0]',Y_train[0], end='\n\n')

    return None

def get_losses(LOSS):
    if LOSS == 'MSE': keras_loss = tf.keras.losses.MeanSquaredError(reduction='none')
    if LOSS == 'MAE': keras_loss = tf.keras.losses.MeanAbsoluteError(reduction='none')
    if LOSS == 'MSLE': keras_loss = tf.keras.losses.MeanSquaredLogarithmicError(reduction='none')
    if LOSS == 'BCE': keras_loss = tf.keras.losses.BinaryCrossentropy(reduction='none')
    if LOSS == 'CCE': keras_loss = tf.keras.losses.CategoricalCrossentropy(reduction='none')
    return keras_loss

def get_activation(ACTIVATION):
    if ACTIVATION == 'sigmoid': activation = sigmoid
    if ACTIVATION == 'tanh': activation = tanh
    if ACTIVATION == 'softmax': activation = softmax
    if ACTIVATION == 'relu': activation = relu
    if ACTIVATION == 'elu': activation = elu
    if ACTIVATION == 'lrelu': activation = lrelu
    if not ACTIVATION: activation = identity
    return activation


def RMSD(K, E, float64_precision):

    K = np.around(K, float64_precision)
    E = np.around(E, float64_precision)

    d = np.around(K - E, float64_precision)
    sd = np.around(np.square(d), float64_precision)
    msd = np.around(np.mean(sd), float64_precision)
    rmsd = np.around(np.sqrt(msd), float64_precision)

    return rmsd


def print_logs(data, extended=True):

    Y_train, epynn_preds, keras_preds, LOSS, keras_loss, epynn_model, keras_end, epynn_end = data

    float64_precision = 16

    keras_preds = np.around(keras_preds, float64_precision)
    epynn_preds = np.around(epynn_preds, float64_precision)

    kloss_keras = keras_loss(Y_train, keras_preds)
    kloss_epynn = keras_loss(Y_train, epynn_preds)
    eloss_keras = epynn_model.training_loss(Y_train, keras_preds)
    eloss_epynn = epynn_model.training_loss(Y_train, epynn_preds)

    if keras_preds.shape[1] == 1:
        acc_keras = (np.around(keras_preds) == Y_train) * 1.
        acc_epynn = (np.around(epynn_preds) == Y_train) * 1.
    else:
        acc_keras = (np.argmax(keras_preds, axis=1) == np.argmax(Y_train, axis=1)) * 1.
        acc_epynn = (np.argmax(epynn_preds, axis=1) == np.argmax(Y_train, axis=1)) * 1.

    kloss_RMSD = RMSD(kloss_keras, kloss_epynn, float64_precision)
    eloss_RMSD = RMSD(eloss_keras, eloss_epynn, float64_precision)
    preds_RMSD = RMSD(keras_preds, epynn_preds, float64_precision)
    acc_RMSD = RMSD(acc_keras, acc_epynn, float64_precision)

    cprint('\nKeras loss function ('+LOSS+') applied on Keras and EpyNN output probs:', attrs=['bold'])
    print(kloss_keras, '(Keras)')
    print(kloss_epynn, '(EpyNN)')
    print(kloss_RMSD, '(RMS)', end='\n\n')

    cprint('\nEpyNN loss function ('+LOSS+') applied on Keras and EpyNN output probs:', attrs=['bold'])
    print(eloss_keras, '(Keras)')
    print(eloss_epynn, '(EpyNN)')
    print(eloss_RMSD, '(RMS)', end='\n\n')

    if extended:
        cprint('Preds from output layer in Keras and EpyNN:', attrs=['bold'])
        print(keras_preds, '(Keras)')
        print(epynn_preds, '(EpyNN)')
        print(preds_RMSD, '(RMS)', end='\n\n')

        cprint('Accuracy from Keras and EpyNN:', attrs=['bold'])
        print(acc_keras, '(Keras)')
        print(acc_epynn, '(EpyNN)')
        print(acc_RMSD, '(RMS)', end='\n\n')

    cprint('Total CPU time:', attrs=['bold'])
    print(keras_end, '(Keras)')
    print(epynn_end, '(EpyNN)', end='\n\n')

    cprint('RMS only:', attrs=['bold'])
    print('{0:<25}'.format(kloss_RMSD), '(1)', 'RMS keras_loss_keras vs keras_loss_epynn')
    print('{0:<25}'.format(eloss_RMSD), '(2)', 'RMS epynn_loss_keras vs epynn_loss_epynn')
    print('{0:<25}'.format(preds_RMSD), '(3)', 'RMS preds_keras vs preds_epynn')
    print('{0:<25}'.format(acc_RMSD), '(4)', 'RMS accuracy_keras vs accuracy_epynn', end='\n\n')

    return kloss_RMSD, preds_RMSD
```
