## Supplementary material for "EpyNN: Educational python for Neural Networks": File S2: README

Reports on:
    - tensorflow==2.3.0 - tensorflow/python/**/*py
    - Number of files and lines of code in tensorflow/python directory.

Content:
    - tensorflow_python.py - Python code to compute data
    - tensorflow_python.xlsx - Results
    - tensorflow/ and tensorflow/python - Empty directories

To reproduce:
    - Copy your local tensorflow/python (e.g. in /home/user/.local/lib/python3.7/site-packages/tensorflow/python for GNU/Linux) to tensorflow/python in the current directory.
    - Run tensorflow_python.py
    - If you get permission errors on files within tensorflow/python/ (current directory), make a dirty fix (e.g. chmod -R 777 tensorflow/python/* for GNU/Linux) and purge once you are done.
