## Supplementary material for "EpyNN: Educational python for Neural Networks": File S2: tensorflow_python.py

```
from pathlib import Path

################################################## LIB
def read_clean_file(f):

    with open(f, 'r') as f:
        raw = f.read().splitlines()

    # Remove blank lines
    raw = [x.strip() for x in raw if x.strip()]

    # Remove block comments
    raw = [x for x in raw if x[0] != '#']

    # Remove docstring
    cleaned = []
    write = True

    for x in raw:

        if x.strip()[0:3] in ['"""', "'''"]:
            # Docstring - START
            write = False

        if write:
            # Out of docstring
            cleaned.append(x)

        if x.strip()[-3:] in ['"""', "'''"]:
            # Docstring - END
            write = True

    return cleaned


################################################## MAIN
# python3.7/site-packages/tensorflow/python
dir_names = ['tensorflow/python/']

for dir_ in dir_names:

    python_files = [str(path) for path in Path(dir_).rglob('*.py')]

    data = {}

    data['data'] = []
    data['paths'] = []
    data['length'] = []

    # For each file
    for path in python_files:
        cleaned = read_clean_file(path)
        data['data'].append(cleaned)
        data['length'].append(len(cleaned))
        data['paths'].append(path)

# CSV output
for path, length in zip(data['paths'], data['length']):
    print(path+','+str(length))


print(len(data['length'])) # Number of files
print(sum(data['length'])) # Number of lines

# Check that there is strictly no line which contains comment markup
data['data'] = [x for x in data['data'] if '"""' in x or "'''" in x or '#' in x]
print(data['data'])
```
